# Western diet increases brain metabolism and adaptive immune responses in a mouse model of amyloidosis

**DOI:** 10.1101/2023.02.15.528645

**Authors:** Marilena Poxleitner, Sabrina H.L. Hoffmann, Georgy Berezhnoy, Tudor Ionescu, Irene Gonzalez-Menendez, Florian C. Maier, Dominik Seyfried, Walter Ehrlichmann, Leticia Quintanilla-Martinez, Andreas M. Schmid, Gerald Reischl, Christoph Trautwein, Andreas Maurer, Bernd J. Pichler, Kristina Herfert, Nicolas Beziere

## Abstract

Diet-induced body weight gain is a growing health problem worldwide, leading to several serious systemic diseases such as diabetes. Because it is often accompanied by a low-grade metabolic inflammation that alters systemic function, dietary changes may also contribute to the progression of neurodegenerative diseases. Here we demonstrate disrupted glucose and fatty acid metabolism and a disrupted plasma metabolome in a mouse model of Alzheimer’s disease following a western diet using a multimodal imaging approach and NMR-based metabolomics. We did not detect glial-dependent neuroinflammation, however using flow cytometry we observed T cell recruitment in the brains of western diet-fed mice. Our study highlights the role of the brain-liver-fat-axis and the adaptive immune system in the disruption of brain homeostasis due to a Western diet.

## Introduction

Overweight and obesity are serious health problems with an increasing prevalence worldwide ^1^. In longitudinal studies of overweight and obese individuals, a changing lifestyle, including less physical activity and poor dietary choices, has shown mid-life obesity and resulting metabolic disorder alterations, e.g. type 2 diabetes, cardiovascular disease, to be a risk factor for developing dementia and cognitive decline decades later ^2–5^. Numerous investigations described the systemic alterations through high-caloric diets like western diets (WDs) ^6–8^. In response to obesity and associated chronic oversupply of fatty acids and sugar, a low-grade chronic inflammation develops, which if persisting over time, leads to a constant release of inflammatory effectors into the periphery ^9–11^. Adipose and hepatic tissues are the main drivers behind this mechanism, and diet-induced severe fatty liver disease has been observed in rodent and human subjects ^6,7,12–14^. Therefore, advancements in understanding the implications of diet-induced obesity for the whole body are an important factor in health research.

Research is still expanding on what is known about the relationship between diet composition, obesity, and the emergence of neurodegenerative disorders and cognitive decline ^15,16^. A current hypothesis is that by the initiation of metabolic and inflammatory processes, such as the proliferation of macrophages in adipose tissue and the release of pro-inflammatory cytokines, immunomodulatory cascades are further activated, eventually leading to neuroinflammation ^17–19^. Triggered by a high-caloric diet, several human and animal studies link the metabolic impact of the diet to the progression of Alzheimer’
ss disease (AD) inducing increased oligomeric Aβ levels and Aβ plaque load in the rodent brain ^18,20–22^. Even more, dietary cues like fatty acids (FAs) and sugars have been shown to modulate central metabolism itself probably increasing the susceptibility to dementia ^23–27^

To date, the molecular mechanisms which connect obesity and AD are not fully understood ^28^. Several ways exist to ensure communication between the CNS and periphery, which allow the CNS to adapt and respond to peripheral cues ^29,30^. In neurodegenerative diseases like Parkinson’
ss disease and AD, emerging evidence indicates that neuroinflammation does not only rely on glial activation, but innate and adaptive immune cells can modulate inflammatory processes in the brain as well ^31–34^. In human brain samples and animal models of AD, immunohistochemical experiments revealed substantial involvement of peripheral innate and adaptive immune system components in the pathogenesis. For example in multiple sclerosis mouse models, self-antigen recognizing T cells have been identified in brains to act as primary drivers of the autoimmune response ^35,36^. Furthermore, infiltration of bone marrow-derived monocytes into CNS was triggered by a high-fat diet ^37^. Study results on the impact of peripheral immune cells on AD pathology are however still not in agreement ^38–42^.

In this study, we show through a multimodal and multiparametric approach that the consumption of a palatable, high-caloric WD during early to mid-life can cause several systemic effects affecting the peripheral and central metabolism. Using the APPPS1 mouse model, a mouse model of early accelerated amyloidosis ^43^, we identified changes in brain metabolism after disease development using different PET tracers and employed MR spectroscopy and metabolomics to investigate systemic alterations. We first investigated changes of cerebral glucose metabolism, using [^18^F]fluorodeoxyglucose ([^18^F]FDG), a well-established PET marker, widely used to investigate cerebral abnormalities. Second, the diet-induced changes of fatty acid metabolism were analyzed using the long-chain fatty acid surrogate 14(R,S)-[^18^F]fluoro-6-thia-heptadecanoic acid ([^18^F]FTHA). Third, to assess diet-induced neuroinflammatory changes in the brain we used the translocator protein (TSPO) tracer [^18^F]GE-180, which is a surrogate marker for neuroinflammation found mainly on activated glia cells after neuronal damage and inflammation ^44–46^. In addition, we performed flow cytometric and metabolic analyses *ex vivo* to investigate the metaflammation profile of the animals in-depth during WD feeding. With PET, we obtained complementary results showing that diet-induced obesity (DIO) and AD had altered brain glucose and fatty acid metabolism, which are independent of the Aβ pathology and microglial activation. Moreover, we identified T cells as an additive factor in the interplay of AD pathology and metaflammation. The imbalance of key plasma metabolites and liver lipids in the periphery, along with the disruption of glucose and fatty acid metabolism in the brain, underscores the importance of a healthy lifestyle and provides further insight into the complex interplay of the brain-liver-fat-axis.

## Methods

### Animals

This study was performed in double transgenic APPPS1-21 mice (B6.Cg-Tg(Thy1-APPSw,Thy1-PSEN1*L166P)21Jckr; APPPS1, n = 21) that co-expressed the human Swedish double mutation APP KM670/671NL and the L166P mutated human PS1 under the control of neuron-specific Thy-1 promotor. This model shows accelerated amyloid deposition at six weeks of age, accompanied in further age by microglial activation ^43^. As controls, wild-type C57BL/6J mice (WT, n = 23) were used. In transgenic and wild-type groups, male and female mice were investigated. ND and WD group littermates were housed in genotype mixed groups in individually ventilated cages with food and water ad libitum in a 12-hour light/dark cycle. All procedures were performed in accordance with German federal regulations on the use and care of experimental animals and approved by the local authorities (Regierungspräsidium Tübingen (R06/21 G)).

### Diet and study design

At the age of 2.1 ± 0.1 months, animals were either fed a western diet (WD, E15721-347, ssniff, Soest, Germany) or a normal rodent diet (ND, V1534-000, ssniff, Soest, Germany) for a total of 24 weeks (Fig. 1a). Compared to the normal rodent diet, the western diet contains higher percentages of fat, sugar and protein which was accompanied by a shifted balance of fatty acids, minerals and trace elements (Suppl. Table 1). To ensure stability of the dietary constituents, WD was entirely replaced once per week. For ND-fed animals, weight was registered starting at 2.9 ± 0.5 months (n = 20; APPPS1 = 9 and WT = 11), whereas for WD-fed animals, registration started at 2.1 ± 0.1 months (n = 24, APPPS1 = 12 and WT = 12). The weighting of animals was conducted weekly. At 7.7 ± 0.4 months, the animals underwent *in vivo* PET and MR imaging over a period of 4 weeks and were sacrificed for further flow cytometric, metabolic, and histologic analyses (Fig. 1a). Notably *in vivo* and *ex vivo* results include individual drop outs of experimental animals due to e.g., technical problems during procedure and/or analyses. In all experiments, mice were randomized and underwent imaging and *ex vivo* experiments in mixed groups of wild-types and transgenics as well as gender.

**Figure 1:**
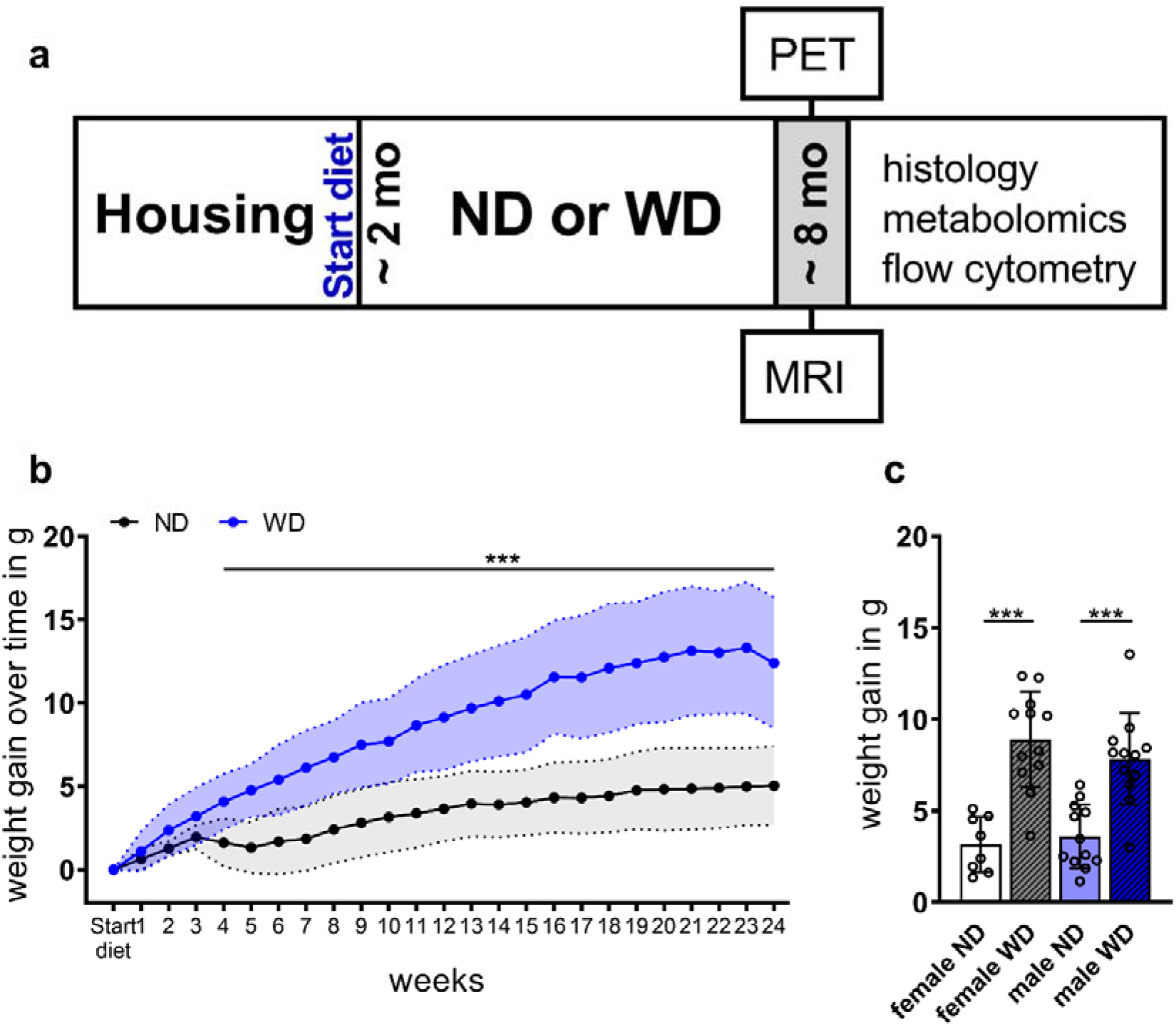
Study design and weight. (a) General study design of in vivo and ex vivo experiments. Western diet (WD) or the normal diet (ND) feeding period started at 2 months age continued over 24 weeks. At ~ 8 months, imaging (PET, MRI), flow cytometry, metabolomics, and histology were performed. (b) Mean weight gain ± SD between ND-fed (black) and WD-fed (blue) animals over the period of 24 weeks starting on the day of the diet change. (c) Mean weight gain between females and males fed an ND (blank white, blue) or WD (striped grey, blue). *** p < 0.001. ND = normal diet; WD = western diet; WD (n=24, male=12, female=12), ND (n=20, male=12, female=8).

### Radiotracer synthesis

Briefly, using the ^18^O(p,n)^18^F nuclear reaction fluorine-18 was produced as [^18^F]fluoride by proton irradiation of [^18^O]H_2_O (Rotem, Leipzig, Germany) at the Tübingen PETtrace cyclotron (GE Healthcare, Uppsala, Sweden).

[^18^F]FDG was synthesized in a TRACERlab MX_FDG_ synthesizer (GE Healthcare, Liège, Belgium) as described previously, using mannose triflate (ABX, Radeberg, Germany) as a precursor ^47^. Quality control was performed according to Ph. Eur. guidelines. Particularly, radiochemical purity, as determined by thin-layer chromatography (TLC), was >95%. Molar radioactivities were > 50 GBq/µmol at the end of synthesis.

[^18^F]FTHA was synthesized using the method from DeGrado ^48^ with modifications on a modified TRACERlab FX_F-N_ synthesizer (GE Healthcare, Münster, Germany). Briefly, 2 µL of the precursor benzyl-14-(R,S)-tosyloxy-6-thiaheptadecanoate (ABX, Germany) in 1 mL of acetonitrile were reacted with a mixture of aceotropically dried [^18^F]fluoride, 15 mg of Kryptofix 2.2.2. and 3.5 mg K_2_CO_3_ at 110°C for 5 min. After hydrolysis with 350 µL of 0.14 N KOH (110 °C, 5 min) 0.3 mL of 6.5 % sulfuric acid was added for neutralization. The product was purified using HPLC (Supelcosil ABZ+; 10 × 250 mm; H_2_O/MeOH 80/20 with 1 % H_3_PO_4_; 5 ml/min; detection: UV 216 nm and NaI(Tl)). The product was obtained in uncorrected yields of 15 ± 5 % (n = 13), corresponding to 9.3 ± 3.3 GBq of isolated [^18^F]FTHA, after irradiations using 35 to 60 µA for 40 to 60 min. Radiochemical purity as determined by TLC was > 90 %. Specific activities were > 50 GBq/µmol at the end of synthesis.

[^18^F]GE-180 was synthesized according to Wickstrøm et al. ^49^ using a FASTlab synthesizer with single-use disposable cassettes (GE Healthcare, Germany) according to manufacturer’s instructions. Quality control was performed via HPLC, yielding the product in chemical purity of > 90% and high molar radioactivity of > 600 GBq/µmol at the end of the synthesis.

### PET imaging

PET studies were performed in C57BL/6J and APPPS1 littermates of each group (ND and WD) over a period of 4 weeks. Animals were anesthetized by using isoflurane (carrier gas 100% oxygen at 1L/min, 5 % for induction, 1.2-1.5 % maintenance) and body temperature was maintained at 37 °C throughout the studies using mouse beds with temperature feedback control (Medres, Cologne, Germany and Jomatik, Tuebingen, Germany). All PET scans were performed using a Inveon dedicated small-animal microPET scanner (Siemens Healthcare, Knoxville (TN), USA), and scans were acquired dynamically for 60 min, immediately followed by a 14 min ^57^Co transmission scan as well as correction of dead time, random and scatter events. Mice were positioned in the center of the field of view and injected intravenously (*i*.*v*.) into a lateral tail vein with 12.0 ± 0.3 MBq [^18^F]FDG (WT-ND n = 10 ; APPPS1-ND n = 7 ; WT-WD n = 7 ; APPPS1-WD n = 8), 14.4 ± 2.3 MBq [^18^F]FTHA (WT-ND n = 8 ; APPPS1-ND n = 8 ; WT-WD n = 8 ; APPPS1-WD n = 7) and 13.5 ± 2.5 MBq [^18^F]GE-180 (WT-ND n = 8 ; APPPS1-ND n = 7 ; WT-WD n = 9 ; APPPS1-WD n = 10) on consecutive days with at least one day of recovery. The mice recovered after each scan on a heating pad in an empty cage, and their health was monitored by the researcher.

### PET image reconstruction and data analysis

List-mode data for all scans were histogramed in 23 frames (8×30 s, 6×60 s, 7×300 s, and 2x 450 s) and reconstructed with two-dimensional ordered subsets expectation maximization (OSEM2D) algorithm with an image zoom of 2 and a 256×256 matrix using Inveon Acquisition Workplace (Siemens Healthcare, USA). Volume-of-interest (VOI) and voxel-wise analyses were performed on reconstructed images using PMOD software v3.2 (PMOD Technologies, Zürich, Switzerland) and statistical parametric mapping SPM 12 (Wellcome Trust Center for Neuroimaging, University College London, United Kingdom). Individual PET images were co-registered to a predefined mouse brain template Mouse-Mirrione-T2 (

PMOD technologies), and a whole-brain VOI as well as a brain-region specific atlas ^50,51^ were applied. The anterior prefrontal cortex area was removed from the cortex VOI to avoid spill-over effects from the harderian glands. The following brain areas were analyzed: cortex (CTX), hippocampus (HIP), cerebellum (CB), and hypothalamus (HYP). Time activity curves (TACS) were extracted and standardized uptake values (SUVs) for each animal were calculated. For comparison of uptake in all four groups, the mean SUV was evaluated between 30-and 60-min post injection (*p*.*i*.). To determine statistical significance one-way ANOVA using multiple comparisons with post hoc Tukey correction was performed using GraphPad Prism 9.0.1 (GraphPad Software LLC, San Diego, USA).

For voxel-wise analysis, PET images were automatically overlaid to Mouse-Mirrione-T2 atlas as reference (PMOD technologies). Differences between groups for each PET tracer were identified using a general linear model (GLM) available in SPM 12. After estimating GLM, statistical parameter maps were generated by interrogating the outcome using contrast vectors. A one-way ANOVA without post hoc correction was applied. Contrasts were compared between groups using no further masking or determined voxel clusters. The significance threshold was set for the tracers individually. Images were prepared using dedicated software (MRIcron, ^52^).

### ^1^H Magnetic resonance spectroscopy of the liver

For magnetic resonance spectroscopy (MRS) on a 7 T BioSpec 70/30 MR scanner (Bruker BioSpin GmbH, Ettlingen, Germany) equipped with a gradient insert, animals were anesthetized using isoflurane (carrier gas oxygen 100% at 1 L/min, 5% for induction, 1.2-1.5% maintenance). Animal body temperature was maintained by placing mice on an MR-compatible water-warmed mouse bed (Jomatik, Tuebingen, Germany). During the whole acquisition, breathing was monitored using a specialized MR breathing pad. Mice were positioned in the center of a ^1^H volume coil with an inner diameter of 86 mm. For correct positioning of liver voxel, an anatomical T2-weighted TurboRARE protocol (TR= 800 ms; TE = 37.63 ms; FOV = 74×32×18; image size = 296×128×72) was acquired. Next, B_0_ map (TR = 30 ms; FOV = 60×60×60 mm^3^, Averages = 1) was acquired. After placing the voxel (3×3×3 mm^3^), avoiding major hepatic blood vessels, the localized shim was acquired resulting in mean shim values of 52.4 ± 13.3 Hz. For spectral acquisition, a stimulated echo acquisition mode (STEAM; TR = 1500 ms; TE = 3 ms; averages = 512) with and without water suppression (VAPOR) was used. All sequences were acquired using Paravision software v6.0.1 (Bruker, Ettlingen, Germany).

Spectral analysis was performed using LC Model analysis software v6.3-1L, (Stephen Provencher, Oakville, ON, Canada; 64). Subsequently, lipid peaks were evaluated according to Ye et al. 2012 ^54^, and the following lipids were extracted: Lip09, Lip13, Lip16, Lip21, Lip23, Lip28, Lip41, Lip43, Lip52, and Lip53. Lipids with a standard deviation (SD) > 20 were excluded ^55^; thus, animal numbers differ between lipids (Suppl. Table 2). Liver fat composition, including lipid mass (LM); fractional lipid mass (fLM), saturated lipid component (SL), fraction of unsaturated lipids (fUL), fraction of saturated lipids (fSL), fraction of polyunsaturated lipid (fPUL), fraction of monounsaturated lipids (fMUL), and mean chain length (MCL) was calculated as described previously ^54^. For LM and fLM lipid peaks Lip13+Lip16 and Lip21+Lip23+Lip28 were united to reduce SD below 20 and hence include all animals into the calculation. Mean values were statistically analyzed using multiple unpaired t-test with post hoc multiple comparison correction using Holm-Sidak method (p value threshold set to α = 0.05) in GraphPad Prism 9.0.1 (GraphPad Software LLC).

### Ex vivo experiments

Following the completion of *in vivo* measurements, the mice were anesthetized by using isoflurane (carrier gas oxygen), and blood was retro-orbitally taken. Then, the mice were sacrificed through asphyxiation with CO_2_ and perfused through the left ventricle with 20 mL of cold PBS, and brain and white adipose tissue (WAT) were removed for *ex vivo* analysis.

### Flow cytometry

Immune cell isolation was performed as described in Hoffmann et al. 2019 ^56^. Briefly, brains and WAT of the abdominal cavity were isolated and chopped into small pieces. Tissue was digested for 45 min in 1 mg/mL Collagenase IV (Sigma Aldrich, St. Lousi, Missouri, USA) in DMEM supplemented with 5% FCS and 10 mM HEPES at 37 °C. Then, digested tissue was washed through a 70 µm mesh cell strainer with 1% FCS in PBS. Brain homogenates were resuspended in 70% percoll (in PBS; GE Healthcare, CA, Illinois, USA) layered under 37% percoll solution topped by a 30% percoll solution in a 15 mL Falcon tube and immediately centrifuged 30 min (acceleration of 2 and deceleration of 1). The immune cells, located between percoll layers 70% and 37% after centrifugation, were isolated and centrifuged for 5 min. The remaining erythrocytes were lysed with 3 mL ACK lysing buffer (Lonza, Basel, Switzerland) for 5 min at room temperature. Following washing, the cell suspension was then pipetted into a 5 mL polystyrene tube via a 40 µm cell strainer snap cap (Corning Inc., Corning, New York, USA). Afterwards, isolated cells from WAT were counted using cell counting chambers (one-way Neubauer counting chambers, C-Chip, Merck, Darmstadt, Germany). Single-cell suspensions were first stained with viability stain (Zombie NIR fixable viability kit, BioLegend, San Diego, California, USA) followed by either BV510-αCD45 (clone: 30-F11), AF700-αB220 (clone: RA3-6B2), BV605-αCD11b (clone: M1/70), BV711-αLy6G (clone: 1A8), BV785-αCD11c (clone: N418), PE/Cy7-αI-A/I-E (clone: M5/114.15.2) and PE-αF4/80 (clone: BM8) or with PE-αCD45.2 (clone 104), FITC-αCD3 (clone 500A2), AF700-αCD8 (clone 53-6.7), BV521-αCD25 (clone PC61), BV510-αCD44 (clone IM7), PE/Cy7-αCD62L (clone MEL-14), BV650-αCD69 (clone H1.2F3), BV785-αCD127 (clone A7R34), BV711-αPD-1 (clone 29F.1A12). All antibodies were purchased from BioLegend (San Diego, CA, USA). The staining took 30 min at 4°C, and cells were afterwards washed three times with PBS, fixed in 0.5% formalin, and analyzed on the BD LSRFortessa flow cytometer (BD Biosciences, Franklin Lakes, New Jersey, USA). Analysis was performed with FlowJo software v10.0.7 (BD Biosciences, USA), and statistical significance was determined using one-way ANOVA corrected for multiple comparisons with post hoc Tukey correction via GraphPad Prism 9.0.1 (GraphPad Software LLC). Gating strategy for both antibody panels are shown in Suppl. Fig 1 and 2. Animal numbers for the organs were for brain (WT-ND n = 11; APPPS1-ND n = 8, WT-WD n = 10, APPPS1-WD n= 9) and for WAT (WT-ND n = 11, APPPS1-ND n = 8, WT-WD n = 10, APPPS1-WD n= 7).

### Metabolomics

For plasma metabolome analysis, blood was collected in an EDTA tube and centrifuged at 4 °C to separate the blood plasma, and aliquots were quenched and snap-frozen with liquid N_2._ A two-phase extraction protocol (polar and lipophilic phases) was applied according to Eggers & Schwudke ^57^. In brief: Blood plasma was transferred to 2 mL AFA glass tubes (Covaris Inc, Woburn, Massachusetts, USA) and mixed with ultra-pure water, tert-butyl methyl ether (MTBE, CAS: 1634-04-4, Sigma-Aldrich Chemie, Taufkirchen, Germany) and methanol. Plasma metabolites were extracted using focused ultrasonication (Covaris Inc, USA) applying the following setup: two treatment cycles, 1^st^: 30 s, Peak Power 125.0, Duty Factor 32.0, Cycles/Burst 400, Avg. Power 40.0. 2^nd^: 30 s, Peak Power 100.0, Duty Factor 30.0, Cycles/Burst 800, Avg. Power 30.0. Temperature range 5.0 to 15.0 °C. Each cycle repeated five times per sample, the total run time per sample was 5 min. Afterwards, the mixture was centrifuged for 5 min, then the polar (water and methanol) phase was decanted. Resulted solution was evaporated to dryness in three hours with a vacuum concentrator (SpeedVac: Preset 2, Thermo Fischer Scientific Inc., Waltham, Massachusetts, USA). Dried pellets of the polar metabolites were resuspended in deuterated phosphate buffer (75 mM Na_2_HPO_4_, 4% NaN_3_, pH = 7.40) with internal standard 3-(trimethylsilyl) propionic-2,2,3,3-d_4_ acid sodium salt (TSP, CAS: 24493-21-8). For maximum dissolution, the Eppendorf cups containing solutions were sonicated and then centrifuged for 5 min aiming to remove any solid residue. The supernatant was transferred into 1.7 mm NMR tubes, then centrifuged for 30 s and subsequently placed in a 96-well rack. Samples were kept cooled (6° C) in the NMR automatic sample handling robot unit - SampleJet (Bruker BioSpin, Karlsruhe, Germany) until the measurement. NMR spectra were recorded by a 14.10 Tesla (600 MHz for proton channel) ultra-shielded NMR spectrometer Avance III HD (Bruker BioSpin, Karlsruhe, Germany) with installed 1.7 mm TXI triple resonance microprobe. NMR measurement routine was performed via a 1D CPMG (Carr-Purcell-Meiboom-Gill) experiment in order to suppress residual background signals from remaining macromolecules like peptides (time domain = 64k points, sweep width = 20 ppm, 512 scans, 1 hour long, temperature 298 K). The recorded free induction decays (FIDs) were Fourier-transformed (FT), and spectra were phase and baseline corrected.

Bruker TopSpin 3.6.1 software was used for spectra acquisition and processing (offset correction, baseline, and phase correction). ChenomX NMR Suite 8.5 Professional (Chenomx Inc., Edmonton, Canada) was used for metabolite annotation and concentration calculation, additionally, internal ChenomX library was included for a resonance frequency of 600 MHz. MetaboAnalyst 5.0 web server (R-based online analysis tool, www.metaboanalyst.ca) was used for metabolite statistical analysis ^58^. Missing values were replaced by a small value (20% of the minimum positive value in the original data). The data was normalized by a reference sample using probabilistic quotient normalization (PQN) ^59^ and scaled using Pareto scaling (mean-centered and divided by the square root of the standard deviation of each variable). Data were analyzed using statistical approaches: one-way ANOVA (analysis of variance), partial least squares discriminant analysis (PLS-DA), and t-testing. Box plot graphical design was performed in GraphPad Prism 9.0.1 (GraphPad Software LLC).

### Histology

Brains were fixed in 4% formalin and paraffin-embedded. For histology, 3-5 µm sections were cut and stained with hematoxylin and eosin (H&E). Immunohistochemistry was performed on an automated immunostainer (Ventana Medical Systems, Inc., Oro Valley, Arizona, USA) according to the company’s protocols for open procedures with slight modifications. The slides were stained with the antibodies CD3 (Clone SP7, DCS Innovative Diagnostik-Systeme GmbH u. Co. KG, Hamburg, Germany), B220 (Clone RA3-6B2, BD Biosciences, Becton, Dickinson and Company, Franklin Lakes, New Jersey, USA), Iba1 (Abcam, Cambridge, UK) and beta-amyloid (Clone Abeta 42, Synaptic Systems, Goettingen, Germany). Appropriate positive and negative controls were used to confirm the adequacy of the staining. All samples were scanned with the Ventana DP200 (Roche, Basel, Switzerland) and processed with the Image Viewer MFC Application. 200x snapshots were taken in all samples in the cortex, hippocampus, thalamus, choroid plexus, and hypothalamus. The number of β-amyloid plaques and the number of B220, CD3, and Iba1 positive cells were determined in those snapshots. Final image preparation was performed with Adobe Photoshop CS6 (final n number: WT-ND n = 2; APPPS1-ND n = 3; WT-WD n = 3; APPPS1-WD n =3).

## Results

### Body weight

To assess the effect of long-term consumption of a WD, WT and APPPS1, animals were fed either ND or a WD starting at the age of 2 months over 24 weeks (Fig. 1a). *In vivo* imaging took place over four weeks and was complemented via flow cytometry, histology and metabolomic experiments (Fig. 1a). Weight monitoring showed a significantly faster weight gain in WD-fed mice compared to ND-fed animals over time (Fig. 1b with a 2.5-fold higher mean weight gain in the WD group (start to end: 8.7 ± 3.8 g) compared to the ND group (start to end: 3.3 ± 1.4 g). Mean weight gain between males and females did not differ (Fig. 1c).

### Liver fat composition and metabolomics

By using ^1^H-magnetic resonance spectroscopy (MRS), we next aimed to assess liver fat composition non-invasively. ^1^H MRS analyses revealed higher lipid fractions with different chain lengths in WD-fed animals (Fig 2a, Suppl. Table 2). We observed a 10-fold higher calculated lipid mass (ND: 0.36 ± 0.15; WD: 3.35 ± 2.00) and fractional lipid mass (ND: 0.48 ± 0.19; WD: 0.88 ± 0.09) in WD-treated mice (Fig. 2b, Suppl. Table 2). Interestingly, among the unsaturated lipid components, only the calculated fraction of polyunsaturated lipids (fPUL) was significantly smaller in WD livers (ND: 0.29 ± 0.10; WD: 0.12 ± 0.10), whereas the saturated lipids (SL and fSL) did not differ between both diets (Suppl. Table 2). Thus, WD leads to a higher accumulation of hepatic lipids. Overall, MR images indicated higher body fat accumulation including abdominal and subcutaneous fat (Fig 2c).

**Figure 2:**
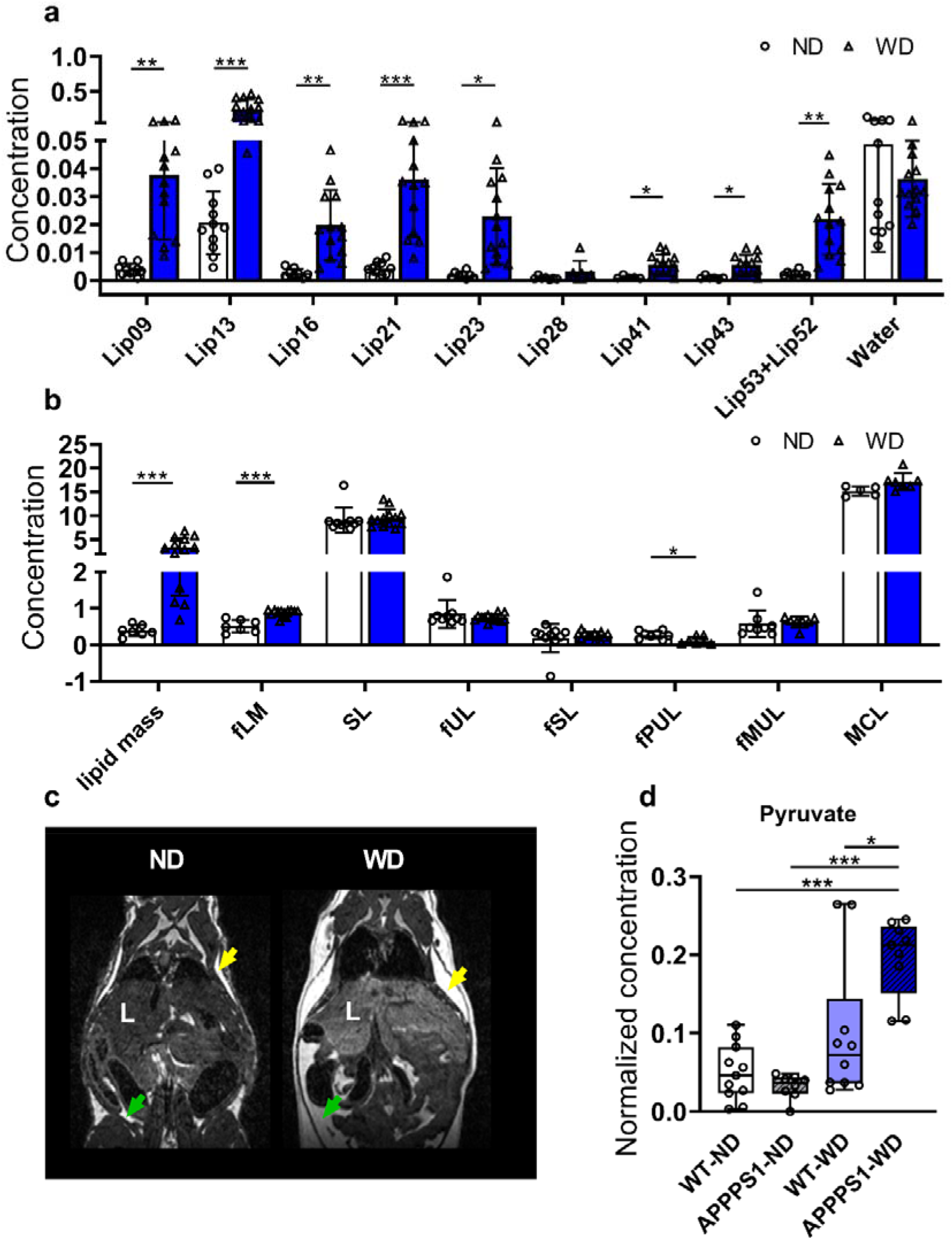
MR-based lipid analysis and metabolomics. ^1^H MRS of hepatic lipid composition and metabolomic results between ND and WD fed animals. (a) single lipids are depicted according to their chemical shift, indicating changes between ND (white, circles)- and WD-fed animals (blue, triangles). (b) Calculated lipid compositions using the single lipid peaks. (c) Exemplary contrast-normalized T2-weighted images illustrating fat depots in ND- and WD-fed mice. Subcutaneous fat marked with yellow arrows; abdominal fat marked with green arrows. (d) Box plot of pyruvate changes between the four mice groups. ^1^H MRS results were analyzed using multiple unpaired t-test with post-hoc multiple comparison correction using Holm-Sidak method (p value threshold set to α = 0.05). fLM = fractional lipid mass; SL = saturated lipid component; fUL = fraction of unsaturated lipids; fSL = fraction of saturated lipids; fPUL = fraction of polyunsaturated lipids; fMUL = fraction of monounsaturated lipids; L = liver; Statistical significance: *p < 0.05, **p < 0.01, ***p < 0.001.

Plasma metabolite profiles were examined by NMR metabolomics. In total, 24 serum metabolites from various metabolite classes, such as amino acids, ketone body – 3-hydroxybutyrate (3-HB), energy metabolites, and short-chain fatty acids were identified. We observed a substantial change for pyruvate in APPPS1-WD animals (Fig 2d; p _ANOVA_ = 8.13E-07; VIP score = 1.4), whereas between the other groups a similar abundance level was detected. All groups, aside from the ND-WT group, developed significantly higher amounts of 3-HB (p _ANOVA_ = 0.00083) and isoleucine (Suppl. Table 3 group A-C; p _ANOVA_ = 0.0016). On the other side, histidine was drastically lowered in the plasma of the APPPS1-WD group (Suppl. Table 3 group C, VIP score = 0.8; p _ANOVA_ = 0.0012). Moreover, considerable changes were identified for glucose (Suppl. Table 3 group B, VIP score = 4.5). Consistently with the higher accumulation of liver fat, animals showed a peripheral misbalance caused by overnutrition. Regression model analysis identified more changes within transgenic and wild-type mice group comparison (Suppl. Table 3 groups A, C). Here, we found lower levels of citrate and succinate – important TCA cycle metabolites, the amino acids phenylalanine and tyrosine, and creatine in transgenic animals.

### Cerebral glucose metabolism

To examine changes in cerebral glucose metabolism, we used [^18^F]FDG-PET. Mean blood glucose values were measured before the imaging and did not differ between groups (Suppl. Fig. 3). Representative images of axially positioned brains of each group showed higher [^18^F]FDG uptake in the APPPS1-WD group. In contrast, no changes between the other groups were observed visually (Fig. 3a). Mean SUV of [^18^F]FDG in the whole-brain displayed significantly higher values in WD-fed APPPS1 mice compared to the other conditions (Fig. 3b). No significant differences in [^18^F]FDG SUV between the other groups could be seen. Further region-based quantification highlighted similar differences between the APPPS1-WD group and the other groups, underlining an overall brain effect. Voxel-wise analysis confirmed a whole-brain effect in the APPPS1-WD group (Fig. 3c). A minor difference in [^18^F]FDG accumulation between WT-ND and WT-WD could be seen in anterior areas, which could not be identified in the prior quantification procedure, indicating a minor effect of WD on [^18^F]FDG in the anterior region of healthy brains. Together with the VOI-based results, [^18^F]FDG revealed hyper glycometabolism in the WD-fed amyloid mouse model.

**Figure 3:**
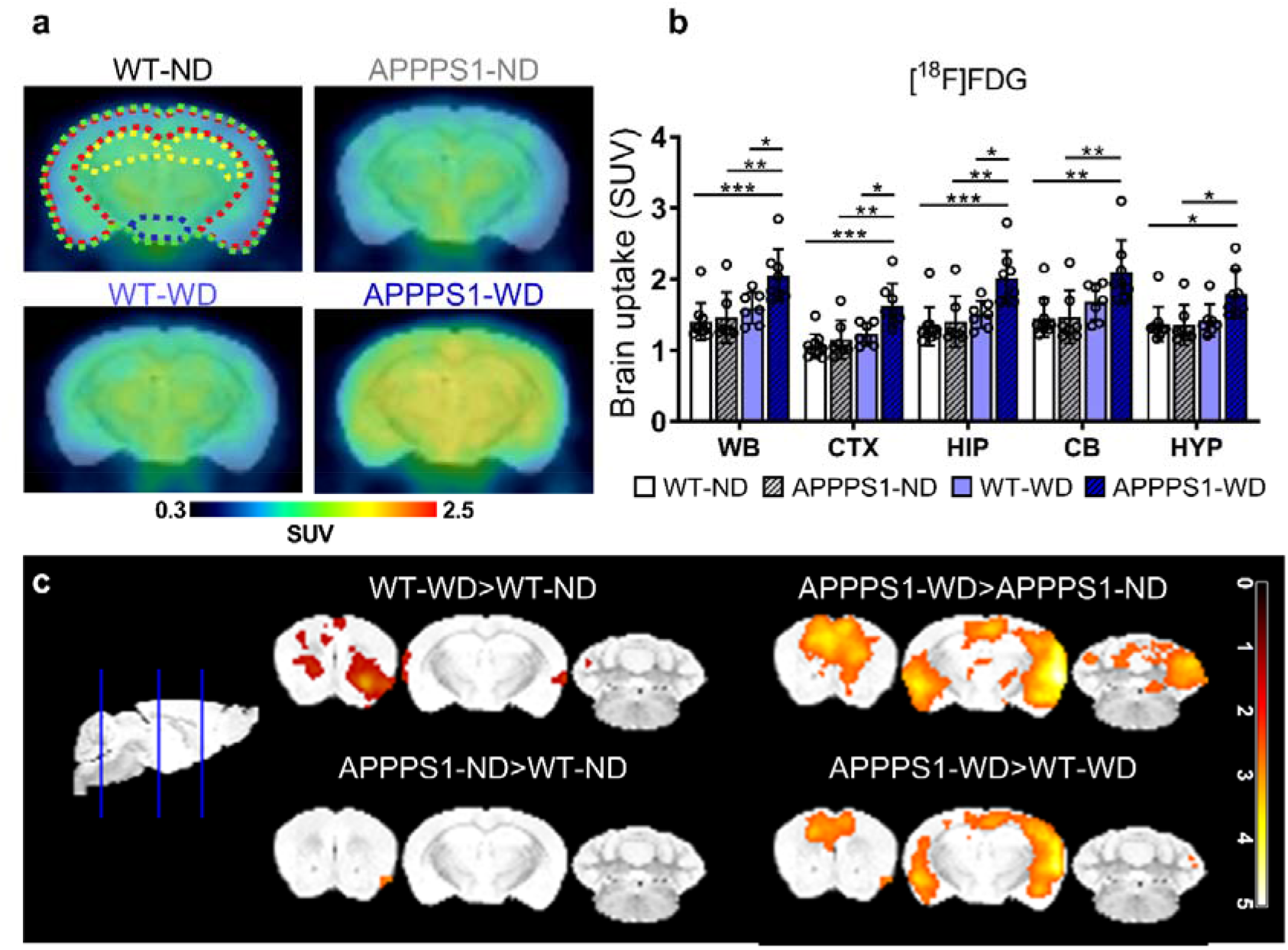
[^18^F]FDG-PET imaging. (a) Comparison of axial brain images of [^18^F]FDG uptake in all four groups indicate a higher uptake in APPPS1-WD mice. Regions are indicated as follows: green = WB; red = CTX; yellow = HIP; blue = HYP. (b) mean SUV (30-60 min *p*.*i*.) in WB, CTX, HIP, CB, and HYP between all groups. (c) T-maps comparing SUVs are shown with threshold p < 0.01. WT-ND n = 10, APPPS1-ND n = 7, WT-WD n = 7, APPPS1-WD n= 8. *p<0.05, **p<0.01, ***p<0.001, post hoc Tukey corrected for multiple comparisons; WB = whole brain; CTX = cortex; HIP = hippocampus; CB = cerebellum; HYP = hypothalamus.

### Cerebral fatty acid metabolism

Next, we investigated the impact of western diet on fatty acid metabolism *in vivo* using the long-chain fatty acid analog [^18^F]FTHA. Axial brain images representing [^18^F]FTHA uptake in all groups displayed a higher brain uptake in WD-fed groups (Fig. 4a). VOI-based whole-brain mean SUVs confirmed a significantly higher [^18^F]FTHA uptake in WD-fed animals, both in WT and APPPS1 animals with no differences between genotypes (Fig. 4b). Segmentation of the brain regions could not highlight any region-specific statistical difference in [^18^F]FTHA accumulation, suggesting a whole-brain effect. Overall, the higher uptake of [^18^F]FTHA in WD-fed mice was independent of the genotype in all observed brain regions. Further data analysis on voxel level showed higher signals in WD-fed animals over ND-fed animals in all observed regions in the brain but did not support identification of more prominent areas (Fig.4c).

**Figure 4:**
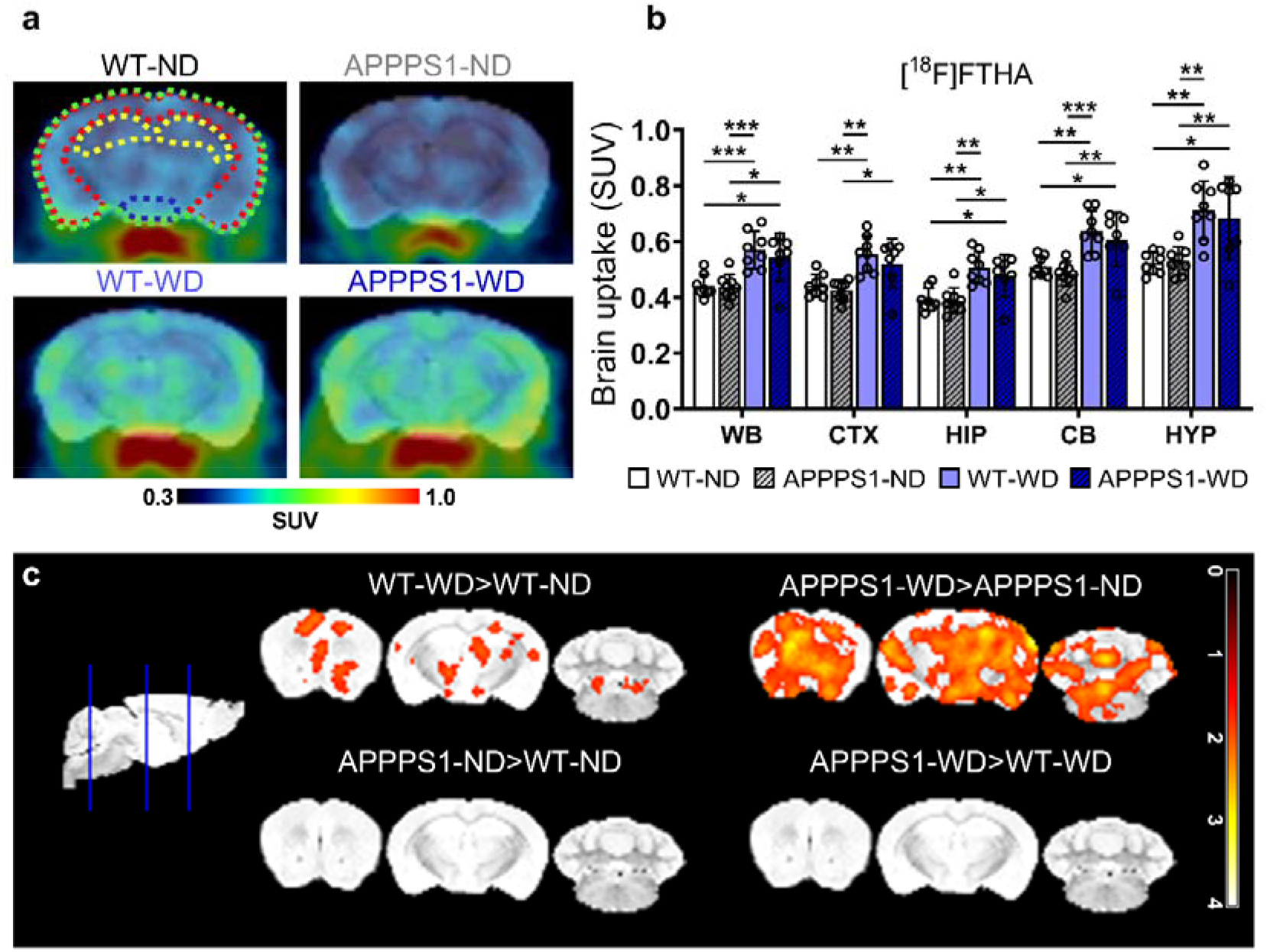
[^18^F]FTHA-PET imaging. (a) Exemplary axial brain images of [^18^F]FTHA uptake display higher uptake in WD-fed mice irrespective of genotype. Regions are indicated as follows: green = WB; red = CTX; yellow = HIP; blue = HYP. (b) Mean SUVs (30-60 min *p*.*i*.) in WB, CTX, HIP, CB and HYP for [^18^F]FTHA. (c) Comparison of voxel-wise analysis. The threshold was set to p < 0.05. WT-ND n = 8, APPPS1-ND n = 8, WT-WD n = 8, APPPS1-WD n= 7. *p<0.05, **p<0.01, ***p<0.001, post hoc Tukey corrected for multiple comparisons; WB = whole brain; CTX = cortex; HIP = hippocampus; CB = cerebellum; HYP = hypothalamus.

### Neuroinflammation

Next, we aimed to investigate the influence of WD on microglia activation in WT and APPPS1 mice using the TSPO-PET tracer [^18^F]GE-180. Whole-brain SUV analysis showed that [^18^F]GE-180 accumulation was significantly elevated in APPPS1 brains, irrespective of the diet (Fig. 5a), pointing to a genotype-dependent effect. Further region-specific analysis pointed at significant radiotracer accumulation differences in the CTX and HIP, regions typically the most affected by amyloidosis (Fig. 5b). Tracer uptake over time revealed a higher injection peak in transgenic animals compared to wild-type animals for cortex, whereas for cerebellum peaks did not differ (Suppl. Fig. 4). Ratios between cortex and cerebellum as well as hippocampus and cerebellum showed significantly higher values in APPPS1 brains than in WT (Suppl. Fig. 5). However, no differences between diets were observed. Further voxel-wise comparison highlighted the main [^18^F]GE-180 accumulation differences between WT and APPPS1 brains to be located in CTX and HIP primarily (Fig. 5c).

**Figure 5:**
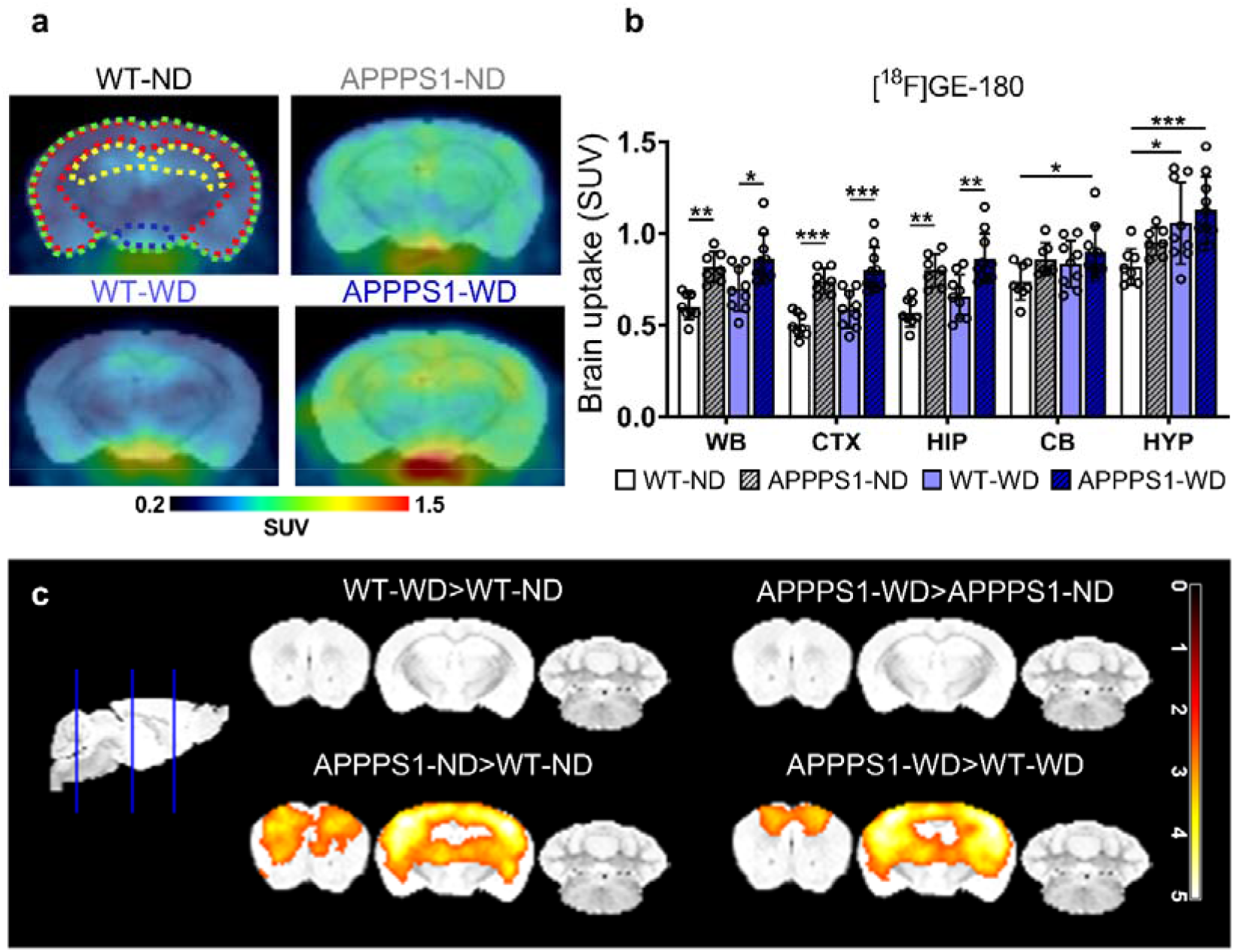
[^18^F]GE-180-PET imaging. (a) Higher uptake of [^18^F]GE-180 in APPPS1 mice compared to WT shown in representative axial brain images. Colored outlines illustrate the analyzed brain regions green = WB; red = CTX; yellow = HIP; blue = HYP. (b) Mean SUVs (30-60 min *p*.*i*.) in WB, CTX, HIP, CB, and HYP for [^18^F]GE-180 in all groups. (c) Representative images of voxel-wise analyzed SUVs are shown with threshold p < 0.01. WT-ND n = 8, APPPS1-ND n = 7, WT-WD n = 9, APPPS1-WD n= 10. *p<0.05, **p<0.01, ***p<0.001; WB = whole brain; CTX = cortex; HIP = hippocampus; CB = cerebellum; HYP = hypothalamus.

### Immune cell presence in the brain

In order to uncover the potential immune changes underlying the brain metabolism and to go beyond glial activation as a marker for brain inflammation as seen by *in vivo* PET, we investigated changes in immune cell infiltration in mice brains. Leukocytes were extracted from brains and sorted between innate and adaptive immune cells by two individual antibody panels, making it possible to check for subtypes. Brains of the APPPS1-WD group had less CD11b^+^Ly6G^+^ neutrophils (Fig 6a; WT-ND vs. APPPSS1-WD p = 0.02; APPPS1-ND vs. APPPS1-WD p = 0.02) than ND brains, no differences could be detected for CD11b^+^ myeloid cells, CD11b^+^F4/80^+^ macrophages, and dendritic cells (CD11c^+^MHCII^+^ DCs). Similar results were observed for the WT-WD group (CD11b^+^Ly6G^+^ neutrophils: APPPS1-ND vs. WT-WD p = 0.05). When investigating T cell infiltration (Fig 6b), the APPPS1-WD group showed a high proportion of CD3^+^ cells, which were elevated compared to WT-ND controls (p= 0.04). Further discrimination between CD8^+^ cytotoxic T cells and CD8^−^ T cells revealed elevated CD8^+^ T cells in APPPS1-WD brains compared to WT groups (APPPS1-WD vs. WT-ND p=0.01; vs. WT-WD p = 0.02). No difference was observed for CD25^+^CD127^−^ regulatory T cells (Tregs). B cell populations did not differ either (Suppl. Fig. 6a). To determine the T cells possible function in the brain, we next investigated the T cell subtypes. Here, WD-fed animals had a higher CD8^−^ T cell effector memory (T_EM_) phenotype, whereas central memory (T_CM_) and naive T cell populations did not change (Fig. 6c). Moreover, these groups had a higher proportion of CD69^+^ lymphocytes, indicating activation, compared to WT-ND controls. In comparison, CD8^+^ T cell subpopulations were elevated only in the APPPS1-WD group (Fig. 6d). Here, a higher percentage of effector memory T cells (T_EM_) compared to the other groups was detected (Fig. 6d; WT-ND versus APPPS1-WD; p=0.02; APPPS1-ND versus APPPS1-WD p= 0.04; WT-WD versus APPPS1-WD p=0.009) and a trend towards higher CD69^+^CD44^+^ activated effector population compared to the WT groups emerged (Fig 6d, WT-ND versus APPPS1-WD p= 0.05; WT-WD versus APPPS1-WD p= 0.04). The immune checkpoint PD1^+^ revealed no differences between any of the groups (Fig 6c, d). While we detected no pronounced infiltration of innate immune cells, these results indicate that WD initiated T cell involvement which displayed an effector state.

**Figure 6:**
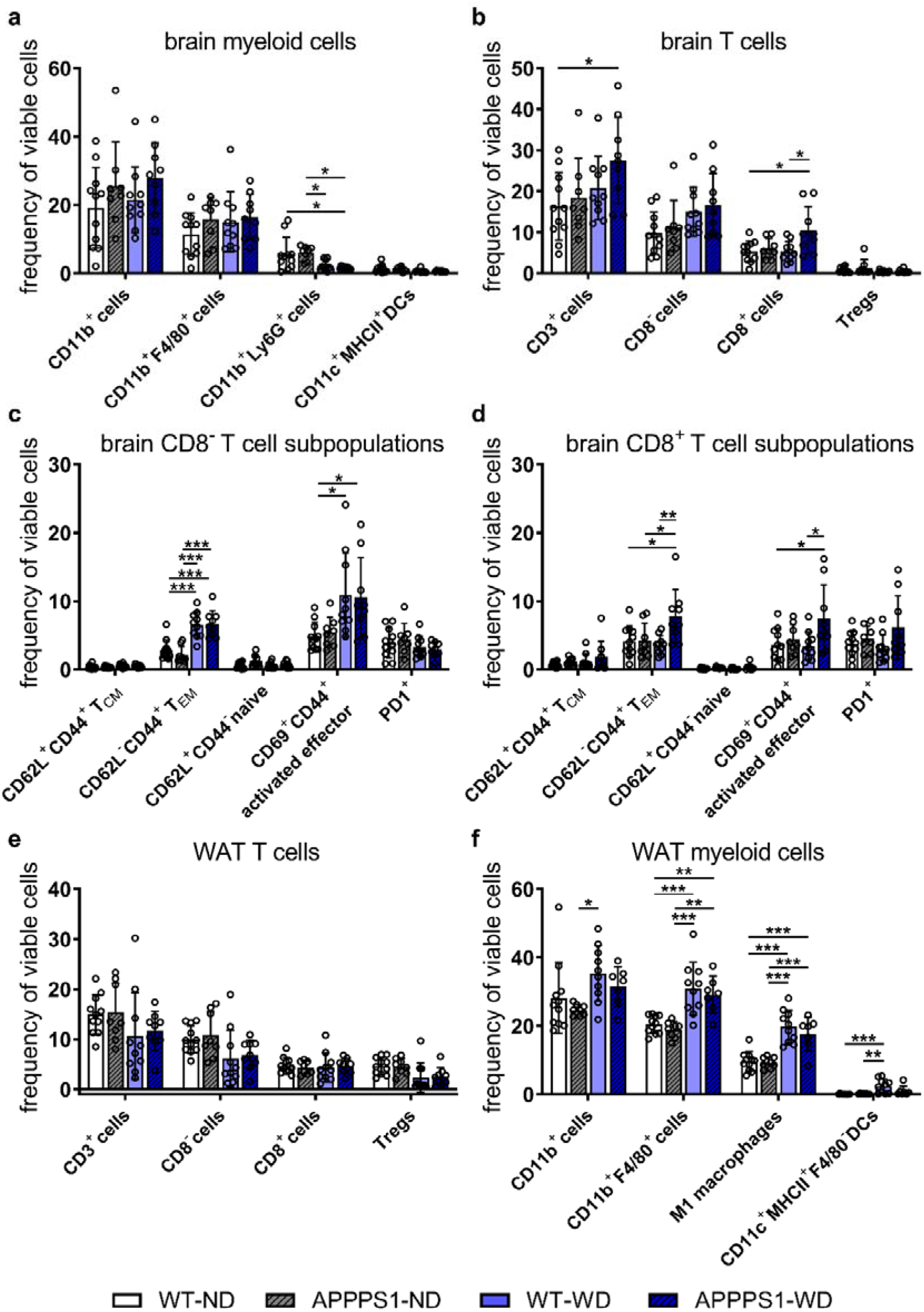
Immune cell analysis. Brain and WAT immune cell population displayed as the frequency of viable cells. (a) Brain myeloid cells show only minor changes. (b) CD3^+^ T cells and CD8^+^ T cells are significantly higher in APPPS1-WD mice compared to non-treated WT animals. (c) CD8^−^ T cell subpopulations reveal higher effector memory T cells (T_EM_) and higher activated effector T cells in WD groups. (d) CD8^+^ T cell populations show higher effector memory and activated effector T cell phenotype, but only in APPPS-WD animals. (e) WAT myeloid cell population display significantly higher CD11b^+^F4/80^+^ macrophages, inflammatory M1 macrophages (CD11b^+^F4/80^+^CD11c^+^) and CD11c^+^MHCII^+^ DCs in WD-fed groups. (f) T cell populations in WAT reveal no differences between groups. Results in mean ± SD; *p<0.05, **p<0.01, ***p<0.001, post hoc Tukey corrected for multiple comparisons; Brain (A-D): WT-ND n = 11, APPPS1-ND n = 8, WT-WD n = 10, APPPS1-WD n= 9. WAT (E-F): WT-ND n = 11, APPPS1-ND n = 8, WT-WD n = 10, APPPS1-WD n= 7. *p<0.05, **p<0.01, ***p<0.001. DC = dendritic cells; T_CM_ = central memory T cells; T_EM_ = effector memory T cells; Tregs = regulatory T cells.

### Immune cell population in white adipose tissue (WAT)

One major hallmark of obesity-induced inflammation is the accumulation and activation of macrophages in adipose tissue ^12^. Therefore, we next investigated changes in T cells and myeloid cells in WAT by flow cytometry (Fig. 6e and f). T cell populations in WAT between groups were not different (Fig. 6e). A substantial elevation of macrophage marker F4/80^+^ was detected in both WD-fed groups. Further examination of the pro-inflammatory macrophage M1 phenotype using CD11c^+ 60^ displayed higher populations in WD groups. The ratio of M1 to M2 F4/80^+^ macrophages was shifted towards a higher M1 portion in WD-fed animals (Suppl. Fig. 7). To ensure that we see dendritic cells (DCs) and not M1 macrophages as all APCs express MHCII and CD11c in WAT, the DC population was additionally gated negative for F4/80. DC populations were elevated in WD groups, in which WT-WD showed the greatest differences. Additionally, B cell populations were significantly higher in WAT of obese compared to lean animals irrespective of their genotype (Suppl. Fig. 6b).

**Figure 7:**
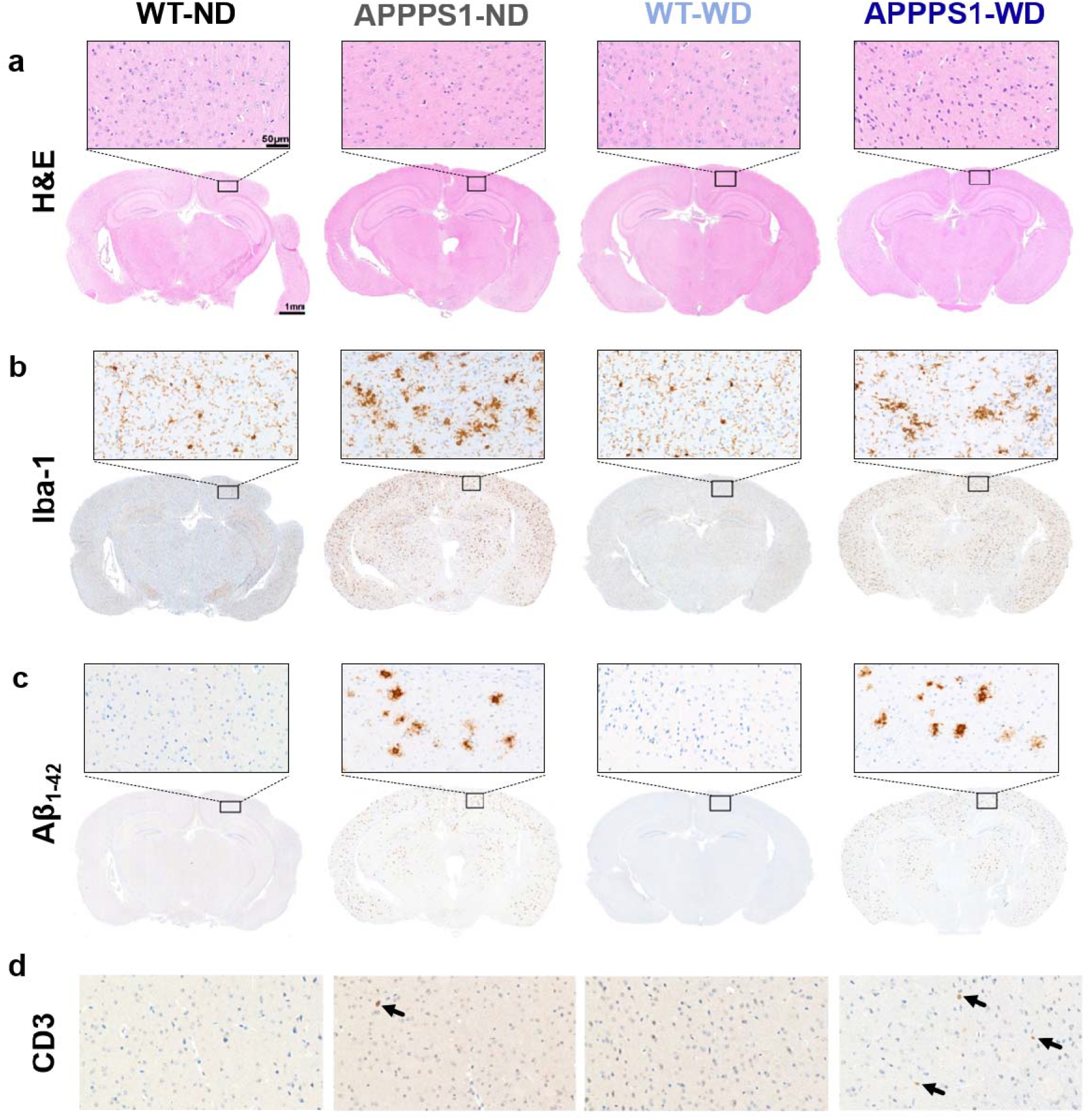
Histological analysis. Histological staining results are depicted in representative images per group per staining (Scale bar 1 mm). Magnifications are depicted for similar cortical areas (Scale bar 50 µm). (a) H&E does not show any anatomical differences between groups. (b) Microglia staining using Iba-1 as marker shows a ramified/resting phenotype in WT brains, whereas activated amoeboid phenotype of microglia in transgenic AD animals was observed, together with a higher number of positive cells. No differences between ND and WD groups were observed. (c) Amyloid plaques stained specifically with Aβ_1-42_ antibody were visible in APPPS1 animals in CTX, HIP, and HYP. WT animals were devoid of specific staining. (d) CD3^+^ staining revealed more T cells in cortical, hippocampal, and hypothalamic regions of APPPS1-WD animals compared to WT animals (black arrows). CTX = cortex; HIP = hippocampus; HYP = hypothalamus. WT-ND n = 2; APPPS1-ND n = 3; WT-WD n = 3; APPPS1-WD n =3.

### Immunohistochemistry

For all analyzed brains, H&E staining revealed no morphological differences between groups (Fig. 7a). To clarify possible differences in microglial activation upon diet and validate our *in vivo* results, brains were analyzed for Iba-1, a microglia marker (Fig. 7b). In transgenic animals, the morphology of microglia changed from a thin and ramified structure of spines to an activated amoeboid structure, confirming an activated phenotype of microglia with higher cell numbers in CTX, HIP, and HYP for transgenic animals. No differences were observed between the diets for the investigated regions, consistent with the observed *in vivo* results. Iba-1 microglia were highly activated in regions of high plaque load (Suppl. Fig 8 black box), whereas they were less activated in regions with few to no plaques (Suppl. Fig 8 red box). Amyloid plaque load in transgenic animals was high, mainly in the CTX and THA, fewer were identified in the HIP, and only a few animals showed plaques in HYP. In WD-fed groups, we could not observe that diet increased plaque load in the investigated brain regions CTX, HIP, and HYP (Fig. 7c). To confirm flow cytometric T cell infiltration in the brain parenchyma, we stained for infiltrating CD3^+^ T cells and found more T cells in AD-WD brains compared to the other groups in CTX as well HIP and HYP (Fig. 7d). Furthermore, for some animals we found a high number near the choroid plexus, the main entry site of peripheral T cells and B cells (Suppl. Fig 9), however intragroup variability was high. In wild-type animals, isolated T cells were observed independent of the diet. Consistent with flow cytometry data, no differences in B cell counts were detected between the brains of all groups (data not shown).

## Discussion

The ageing of the general population in western countries is accompanied by an increase in the prevalence of dementia, and it is suspected that the increase in overweight and obesity exacerbate this challenge in public health. The suspected underlying link involves a general chronic inflammatory state of the patient called metaflammation, but further work is required to understand the intricacies governing cerebral metabolic disruption and unbalanced diets. In this work, we investigated such potential interaction first by using non-invasive imaging techniques to identify molecular and metabolic dysfunctions in different organs *in vivo*. In our experiments involving wild-type mice and a murine amyloidosis model, a surrogate for AD progression, mice were fed a western diet upon manifestation of amyloid pathology, covering the preclinical early to mid-life period when pathophysiological changes can already be detected ^61^. We, therefore, used an amyloidosis mouse model which is well described ^43,62^ and represents an un-modifiable AD risk factor. Importantly, our model does not include age-related effects on the brain and periphery induced by the so-called inflammaging, a phenomenon proposed to be a low-grade systemic inflammatory process ^63^ also favoring age-related diseases like AD ^64^. We chose for this study a western diet which is known to mimic the nutrition of western countries with high fat and high sugar content together with simple carbohydrates and a shifted fat composition towards saturated fatty acids ^65^. This led to a significant weight gain increase when fed over six months in both, male and female mice.

As the liver is one of the organs heavily affected by a high-caloric diet leading to systemic disruption and metabolic imbalance, we wanted to monitor the grade of fatty liver syndrome in WD-fed animals. Via non-invasive proton magnet resonance spectroscopy (^1^H MRS), we detected markedly elevated lipid levels in the livers of WD-exposed mice. Similar results could also be detected in the livers of patients already after a 2-week HFD ^66. 1^H MRS revealed significantly higher lipid mass and fractional lipid mass together with lower levels of polyunsaturated lipids in WD compared to ND livers, which mirrors the evolution of liver composition in rats fed with high-fat diet ^67^. In another study, comparing leptin-deficient ob/ob mice to controls, a decrease in polyunsaturated lipids and an increase in saturated lipids was measured in this obesity-only model excluding dietary impact ^68^. However, we could not detect differences in saturated lipids between the diets. Even though some assume a homogenous hepatic fat distribution ^69^, others report heterogeneous hepatic fat distribution following HFD ^70,71^, making it likely that the spectra of the positioned single voxel do not reflect the whole liver condition. The changes in single hepatic lipid peaks and lipid fractions after WD endorse the impact of the diet on liver fat accumulation, even after 6 months of WD. Metabolite analysis of plasma revealed significant differences for pyruvate, 3-hydroxybutyrate (3-HB), histidine, and isoleucine. 3-Hydroxybutyrate has been shown to function in rodents as an anti-AD drug ^72^ and neuroprotective agent ^73^. In our study, the levels are found to be highest in ND-fed transgenic animals, but we found high levels in WD-fed animals too. Moreover, levels decline in WD-fed APPPS1 animals, which we found to have brain glucose hypermetabolism. Together with a contradicting study, which showed3-HB to be high in 3xTg animals but low in HFD-fed animals and was associated with glucose metabolism compensation ^74^., 3-HB might be an important plasma marker to indicate brain glucose metabolism changes. Administration of histidine to mice brains has been postulated to alleviate chronic effects of hypoperfusion by, among other, improving BBB integrity ^75^. Even further, histidine application is associated with a neuroprotective role in AD ^76^. Here, the lower levels only in APPPS1-WD animals might indicate the acceleration of detrimental processes in this group and in combination with other plasma markers (3-HB, pyruvate) could be used to specify central metabolic disruptions further.

In our *in vivo* imaging approach, we investigated diet-induced effects on brain metabolism. By using glucose and long-chain fatty acid surrogates, we could monitor brain metabolism alterations induced by diet in wild-types and APPPS1 mice. [^18^F]FDG brain uptake was higher in WD-fed APPPS1 mice compared to the other conditions, indicating that WD leads to hypermetabolism in AD transgenic animals. Analysis of the voxel level confirmed a whole-brain effect in APPPS1-WD animals. As studies have already shown a positive correlation between the [^18^F]FDG and [^18^F]GE-180 signal in aging wild-type mice assuming higher glucose demand due to higher glial activation ^77^, comparison of both tracers revealed in our study no higher glial-dependent neuroinflammation in APPPS1-WD group compared to the other groups. This was confirmed by Iba-1 staining in brain tissue. Although obtained using a different diet and model, these results can be compared to studies done using mice infused with human Aβ_42_ while fed an HFD over three months ^78^. The authors could show a [^18^F]FDG hypermetabolism when diet and Aβ were combined and saw no association between TSPO signal and glucose uptake, assuming that gliosis is not the only player in diet-induced neuroinflammation. Longitudinal assessment of [^18^F]FDG brain uptake in the same transgenic model has been shown to not differ from controls in mid-age, similar to our results, but decrease with advanced age ^79^, assuming that the consumption of a WD can initiate hypermetabolism in this amyloid model. Interestingly, Ashraf et al. claimed that the hypermetabolic phase they observed in patients with mild cognitive impairment (MCI) might reflect a compensatory neuroplastic mechanism of neurons, which, when overstimulated, could exhaust, thereby accelerating the degenerative process ^80^. Thus, the high [^18^F]FDG uptake that we see in the APPPS1-WD group may represent a transient process of a neuronal compensatory response that eventually leads to neuronal death and cognitive decline as a consequence of diet-induced obesity (DIO) and/or dietary components. We clearly can emphasize a central as well as a systemic disruption in glucose metabolism as metabolomic analysis revealed elevated pyruvate plasma levels in the WD-fed transgenic animals complementing the PET results. In addition, metabolomic results revealed a higher VIP (Variable Importance in Projection) score for glucose; however not significant in the ANOVA analysis.

Many studies could show that direct or indirect (via diet) supplementation of peripheral fatty acids can activate inflammatory cascades in the brain e.g., via TLRs, and therefore induce inflammatory processes ^81^. We used the long-chain fatty acid tracer [^18^F]FTHA to determine fatty acid metabolism when continuous delivery of fatty acids is given. Brain uptake revealed indeed a diet-dependent higher fatty acid metabolism, which was independent of the genotype. In human and pig brains, [^18^F]FTHA has been shown to cross the BBB and to represent central fatty acid oxidation ^82,83^. Moreover, in patients with metabolic syndrome, [^18^F]FTHA brain uptake was higher, similar to our results. Studies propose that high levels of saturated FA could lead to the activation of microglia and astrocytes ^84,85^, so we compared [^18^F]GE-180 signals to the [^18^F]FTHA signals with both VOI-based and voxel-wise analysis, but could not detect overlapping regions. However, no further discrimination between normal and diseased brains was found in this model using [^18^F]FTHA.

Neuroinflammation was assessed by imaging using the TSPO tracer [^18^F]GE-180, which revealed higher uptake in pathology-rich regions of the transgenic brain, correlating with results obtained from other studies using other AD models ^46,86,87^. In contrast with other studies showing a higher glial activity after energy-rich diets (for a comprehensive overview see ^15^), we could not demonstrate any diet-dependent variations in this model. It is, however, also possible that the feeding duration or composition of our WD might not initiate higher glial inflammation. For instance, in a study that used two high-caloric diets in the same experimental set-up, only the diet with high lard content (60% fat) led to higher microglial activation, whereas the WD (40% fat) did not ^88^. Furthermore, different durations of HFD seem to employ region-specific inflammatory processes in the cortex and the cerebellum of mice ^89^.

Systemic and central alterations caused by the chronic consumption of a high-caloric diet or the AD pathology per se might affect tracer uptake into the brain. By determining regional cerebral blood flow (rCBF) no changes in perfusion were reported for our AD model ^90^ and in mice fed an WD for 12 weeks ^91^, even after HFD for six months, a lower perfusion was measured ^92^. However, other studies report no increased permeability ^89,92^. In DIO rats fed a WD, an elevated BBB permeability was not observed earlier than 90 days, suggesting a gradual BBB breakdown ^93^. In our study, the western diet might act as an additive factor for BBB permeability in amyloid-prone animals by disturbing brain metabolic balance. However, further investigations need to clarify this hypothesis.

The neuroinflammatory concept is constantly under revision and extensive work in this field is ongoing ^94^, supporting evidence in addition to the initial amyloid cascade hypothesis, that systemic alterations act as neuroinflammatory drivers by activating inflammatory processes e.g. by immune cell infiltration and activation. To determine immune cell involvement in the brain, we examined infiltration of innate immune cells as they have been shown to invade brains after HFD treatment ^37^ as well as in AD-prone mice models ^40^. Except for lower Ly6G^+^ populations, no higher infiltration of innate immune cells was detected, but we could find a higher number of CD3^+^ cells in the group of APPPS1-WD whose infiltration we validated via histological staining. Further discrimination into CD3^+^CD8^−^ T helper cells and CD3^+^CD8^+^ cytotoxic T cells revealed significantly higher cytotoxic T cells in the WD-fed transgenic animals compared to the wild-types. In patients of advanced stages of AD, a study could also find a higher CD3^+^ T cell population in brains, which were CD8 positive ^95^. We could not identify T cells near plaques in our model, which is in line with other reports that have shown T cells to be present in mouse models of AD, but could not see interaction with the plaques or tau pathology ^96,97^. The role T cells play in neurodegenerative disease and which mechanisms the infiltrating T cells initiate once they reside in the parenchyma is still discussed. Several studies point towards a neuro-protective role in AD mouse models ^42^, while others report detrimental effects ^41^. Further discrimination of T cell subtypes in our study, could show a polarization towards an effector memory or activated effector phenotype of both CD8^−^ and CD8^+^ T cells in the WD group. The chronic metabolic inflammation in organs like adipose tissue, which display a constant pro-inflammatory burden for the body, might facilitate activation of naïve T cells in immune compartments of the periphery before they enter the CNS ^98,99^. Whether the T cells were polarized by signals from the periphery or via CNS internal signals and to which extent they disturb CNS homeostasis and accelerate inflammatory processes needs to be further clarified.

The investigation of the metaflammatory condition in white adipose tissue (WAT) in the periphery has proven to correspond to studies describing macrophage recruitment and polarization towards pro-inflammatory status in inflamed WAT ^100^. Although we found higher B cell populations in WD-WAT, which contribute to systemic inflammation by modulating T cells and macrophages ^101–103^, no significant changes in T cell populations were detected in our model. Overall, results clearly show disruption of innate immune cell infiltrates in WAT of WD-fed animals with minor changes in T cell populations. To our knowledge, we were the first to compare WAT immune cells of APPPS1 and wild-types, which showed no differences for both diets. Further experiments to distinguish T cell phenotypes would be helpful to examine the impact of the WD on the T cells in WAT.

We are aware that our study includes limitations, which are discussed in the following paragraph. For the *in vivo* studies, we chose to compare the SUV to correct for the significant weight changes between diet groups. We chose to not compare SUV ratios due to the lack of an adequate reference region in our project. Mostly the cerebellum is used as a pseudo-reference region, but its uptake changed significantly between WT under the control diet and AD mice under WD for [^18^F]FDG and [^18^F]FTHA. That the cerebellum is affected by the diet has already been observed in another study ^89^. When interpreting [^18^F]FDG results from different studies, several factors should be considered. The chosen AD models seem to have a significant impact on the outcome of studies looking at the brain metabolism with a decreased ^79,104^, increased ^86,105^, or no different brain [^18^F]FDG uptake ^106^. It is important to note that the TSPO tracer, [^18^F]GE-180 has been the subject of an extensive debate as several studies have shown that the tracer has only very low to no brain uptake and that this is only sufficiently high when the BBB is disrupted due to the pathology ^107^. Our results show higher brain uptake of [^18^F]GE-180 in transgenic mice compared to wild-types for the pathology-rich cortex, which could be related to an altered BBB in WD-fed APPPS1 mice as discussed previously. Unfortunately, other second generation TSPO tracers have several problems, such as mixed ligand binding affinity due to gene polymorphism and thus the development of alternative inflammatory tracers is of enormous importance for future studies in neurological disorders ^108^.

In this study, we propose that in AD-prone brains, further mechanisms are triggered by a WD beyond the classical glial neuroinflammation. Moreover, we encourage further studies to examine the relation of T cells and brain glucose metabolism in AD, as both were elevated in the amyloidosis model after the WD.

## Acknowledgements

We thank Phillip Knopf, Vera Jörke, Simon Freisinger, Marta Vuozzo, Laura Kübler, Carsten Calaminus, Max Zimmermann and Sabrina Buss for their valuable support in this study. Mathias Jucker and Synovo GmbH for providing the APPPS1-21 transgenic model. Natalie Hermann, Maren Harant, Linda Schramm, Dennis Haupt, Sandro Aidone and Miriam Owzcorz for excellent technical support. Barbara Schörg, Dominik Sonanini, Manfred Kneilling for provision of the flow cytometry T cell antibody panel. Bruker BioSpin GmbH for the support in the NMR-related part of the study. The Facs Core Facility of the Medical Clinic in Tuebingen for the support of the flow cytometry measurements. This study was supported by the Werner Siemens Foundation to B.J.P., and the fortune grant to F.C.M.

## Contributions

M.P. wrote the manuscript and performed in vivo imaging, ex vivo flow cytometry and respective analyses. S.L.H. contributed in in vivo, ex vivo experiments and analyses. G.S.B & C.T. performed metabolomics and analyses. T.I. wrote the code for the voxel-wise analysis and performed pre-processing. I.G-M. and L.Q-F. performed histological staining and analyses. F.C.M. developed the concept. D.S., W.E., G.R., A.M. did radiotracer synthesis and validation. A.M.S. contributed the 1H spectroscopy experiments and analyses. B.J.P. supervised and provided laboratory and equipment. K.H. & N.B. supervised and revised and edited the manuscript. All authors revised manuscript and all agree with its content.

