## Supplementary Materials for "Western diet increases brain metabolism and adaptive immune responses in a mouse model of amyloidosis"

Supplemental Table 1

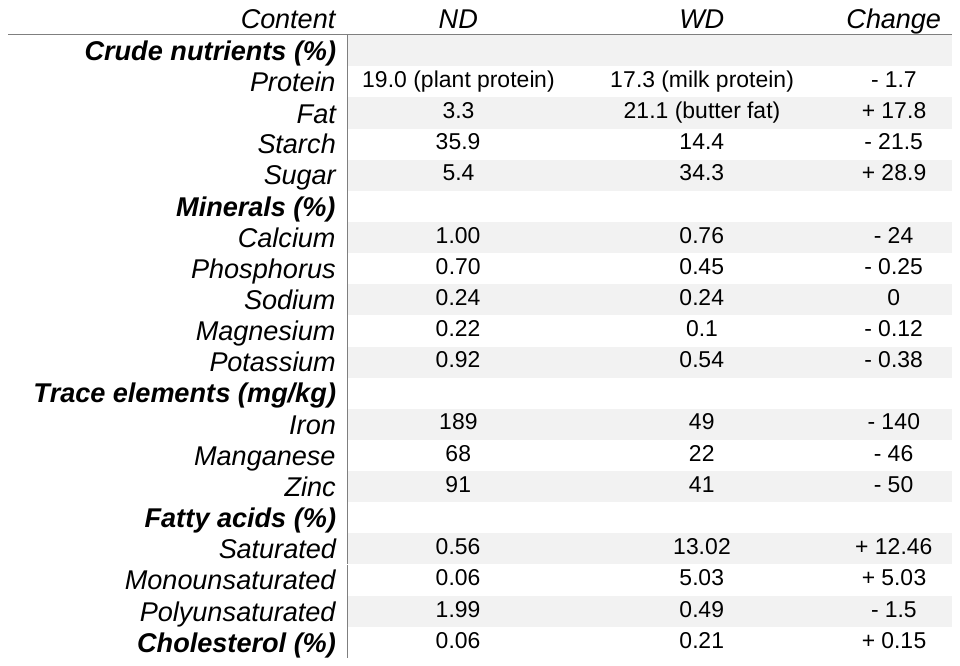

Supplemental Table 1:

Comparison between content of the normal diet and the western diet. Changes indicate higher or lower portions in WD compared to ND for each compound.

Supplemental Table 2

| Lipids | ND | | | WD | | |
| --- | --- | --- | --- | --- | --- | --- |
|  | **mean** | **SD** | **n** | **mean** | **SD** | **n** |
| Lip09 | 0,004 | 0,002 | 10 | 0,038 | 0,023 | 13 |
| Lip13 | 0,021 | 0,011 | 11 | 0,240 | 0,145 | 13 |
| Lip16 | 0,003 | 0,002 | 8 | 0,020 | 0,013 | 13 |
| Lip21 | 0,004 | 0,002 | 10 | 0,036 | 0,021 | 13 |
| Lip23 | 0,002 | 0,001 | 8 | 0,023 | 0,017 | 13 |
| Lip28 | 0,001 | 0,001 | 7 | 0,003 | 0,004 | 7 |
| Lip41 | 0,001 | 0,001 | 5 | 0,006 | 0,004 | 13 |
| Lip43 | 0,001 | 0,001 | 6 | 0,006 | 0,004 | 13 |
| Lip53+Lip52 | 0,003 | 0,001 | 8 | 0,022 | 0,013 | 13 |
| Water | 0,049 | 0,038 | 11 | 0,036 | 0,014 | 13 |
| lipid mass | 0,357 | 0,146 | 9 | 3,347 | 2,004 | 13 |
| fLM | 0,484 | 0,187 | 9 | 0,8771 | 0,093 | 13 |
| SL | 8,315 | 0,542 | 9 | 9,510 | 1,857 | 13 |
| fUL | 0,852 | 0,384 | 9 | 0,7426 | 0,098 | 13 |
| fSL | 0,273 | 0,083 | 8 | 0,258 | 0,098 | 13 |
| fPUL | 0,295 | 0,010 | 7 | 0,116 | 0,105 | 7 |
| fMUL | 0,563 | 0,395 | 7 | 0,625 | 0,150 | 7 |
| MCL | 15,493 | 1,562 | 9 | 16,684 | 2,412 | 13 |

Supplemental Table 2:

Lipids of different chain lengths and lipid compositions are listed. Mean values, standard deviation (SD) and animal numbers (n) for ND and WD are depicted.

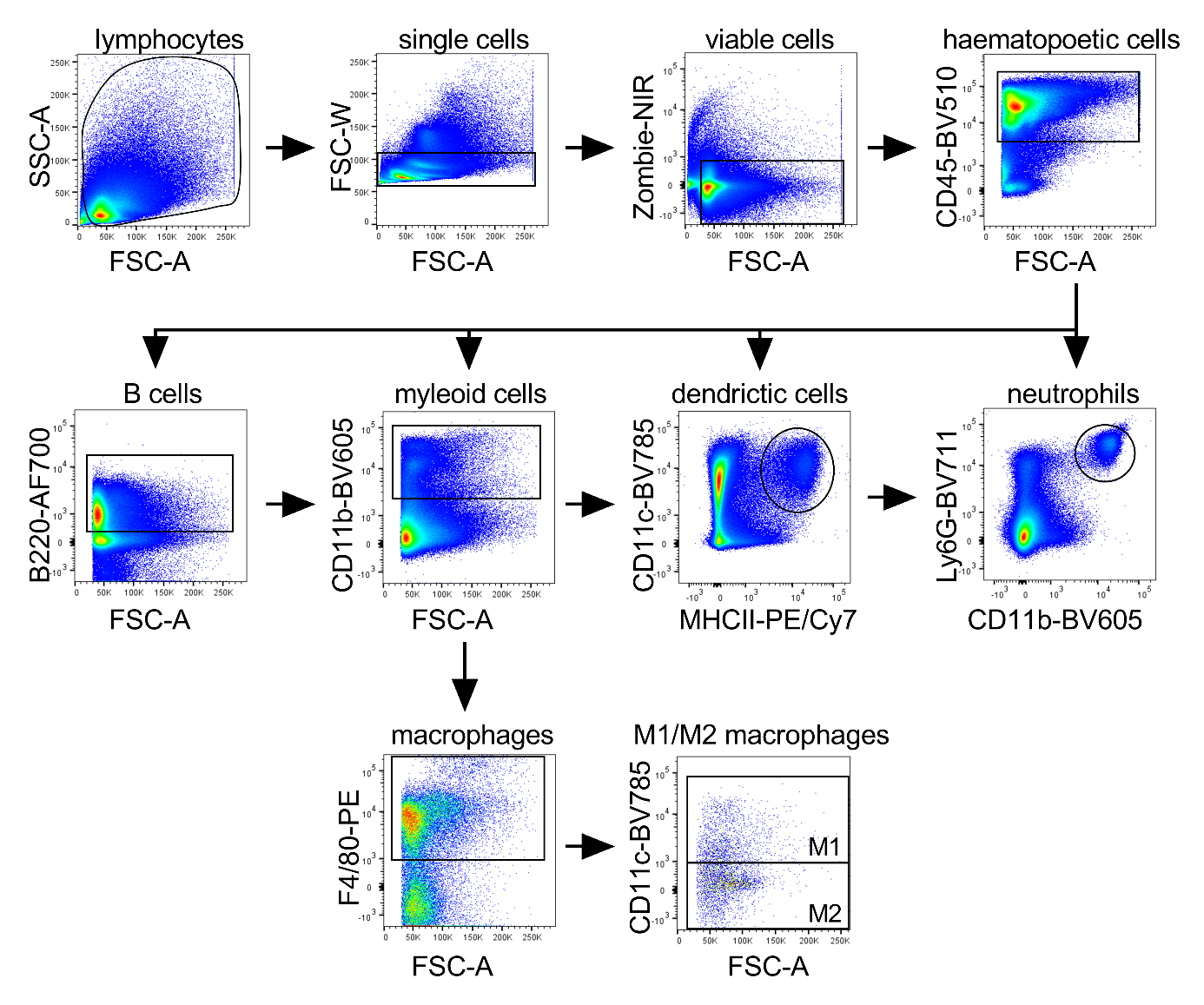
Supplemental Fig.1

Supplemental Figure 1:

Gating strategy for myeloid flow cytometry antibody panel. For each immune cell population the respective antibodies are displayed on the Y- and X-axis. Gating is shown as black outline within the plots.

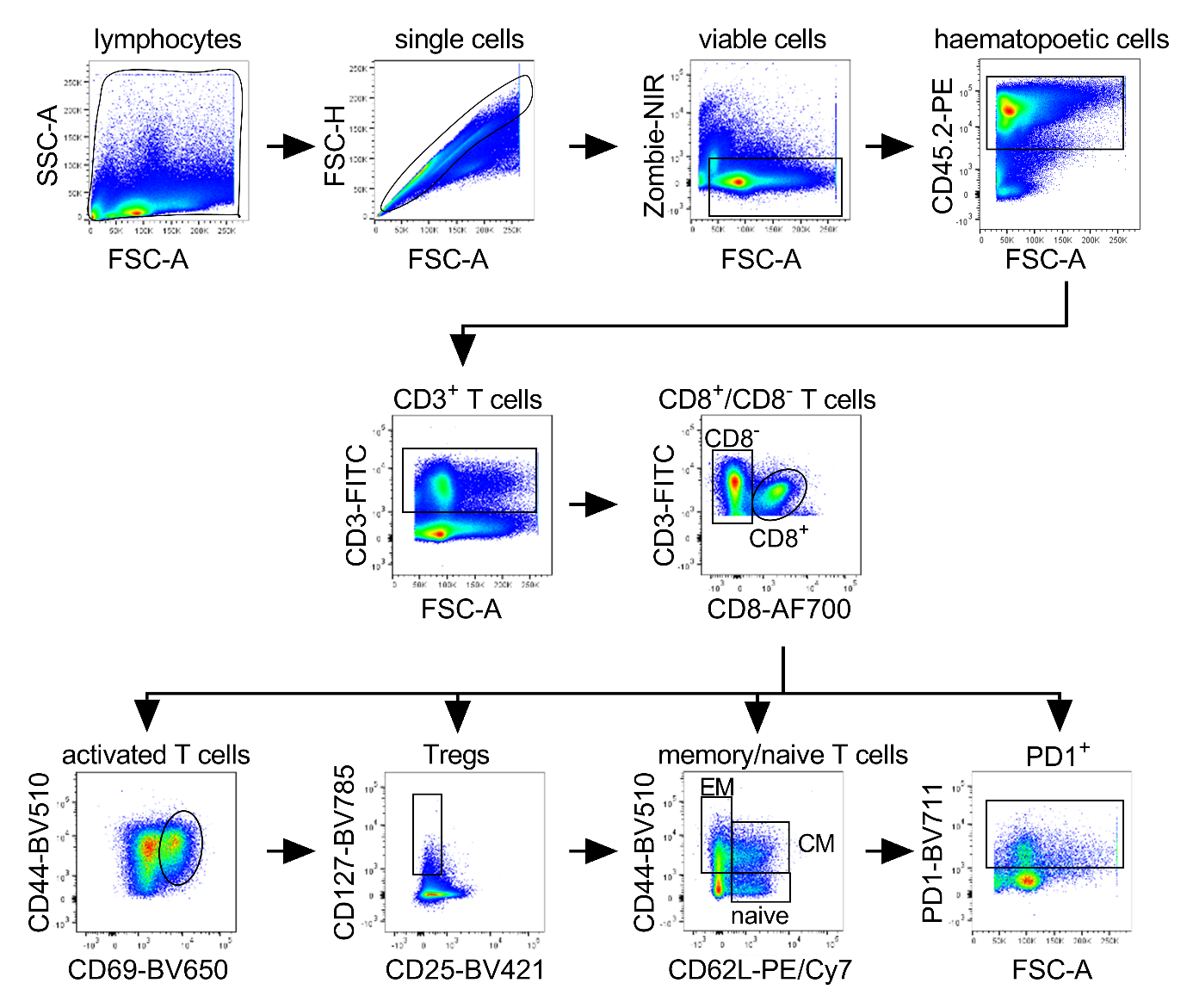
Supplemental Fig. 2

Supplemental Figure 2:

Gating strategy for T cell flow cytometry antibody panel. For each immune cell population the respective antibodies are displayed on the Y- and X-axis. Gating is shown as black outline within the plots. Gating for Tregs applies only for CD8^-^ T cell populations.

Supplemental Table 3

| Description | Metabolite | APPPS1-ND / WT-ND | | | WT-WD / WT-ND | | | APPPS1-WD / WT-ND | | |
| --- | --- | --- | --- | --- | --- | --- | --- | --- | --- | --- |
|  |  | **A** | VIP | | **B** | VIP | | **C** | VIP | |
| **Ketone body** | **3-Hydroxybutyrate** | **↑***** | | **1.7** | **↑**** | | **1.0** | **↑*** | | **0.5** |
| **Short chain fatty acids** | Acetate | ↑ | | 0.8 | ↓ | | 0.1 | **↑** | | 0.1 |
|  | Formate | ↓ | | 1.8 | ↓ | | 0.5 | **↑** | | 0.1 |
|  | Isobutyrate | ↑ | | 0.2 | **↑** | | 0.1 | **↑** | | 0.1 |
| **Amino acids** | Alanine | ≈ | | < 0.1 | ↓ | | 0.2 | **↑** | | 1.1 |
|  | Glutamine | ↑ | | 0.9 | ↓ | | 0.3 | **↑** | | 0.2 |
|  | Glycine | ↑ | | 0.1 | ↓ | | 0.3 | **↑** | | 0.7 |
|  | **Histidine** | ↓ | | 0.5 | ↓ | | 0.1 | **↓***** | | **0.8** |
|  | **Isoleucine** | **↑**** | | **1.0** | **↑**** | | **0.7** | **↑***** | | **0.6** |
|  | Leucine | ↓ | | 0.3 | **↑** | | 0.2 | ↓ | | 0.1 |
|  | Lysine | ↑ | | 0.5 | ↓ | | 0.6 | ↓ | | 0.7 |
|  | Methionine | ↓ | | 0.1 | ↓ | | 0.1 | ≈ | | < 0.1 |
|  | Phenylalanine | ↓ | | 0.5 | **↑** | | 0.1 | ↓ | | 0.3 |
|  | Proline | ↑ | | 0.4 | ↓ | | 0.3 | ↓ | | 0.2 |
|  | Tyrosine | ↓ | | 0.6 | **↑** | | 0.1 | ↓ | | 0.5 |
|  | Valine | ↓ | | 0.2 | ↓ | | 0.1 | ↓ | | 0.1 |
| **Krebs cycle (TCA)** | Citrate | ↓ | | 0.9 | ↓ | | 0.1 | ↓ | | 0.8 |
|  | Fumarate | ↓ | | 0.3 | ↓ | | 0.3 | ↓ | | 0.1 |
|  | Succinate | ↓ | | 1.5 | ↓ | | 0.9 | ↓ | | 1.0 |
| **Creatine metabolism** | Creatine | ↓ | | 0.3 | ↓ | | 0.2 | ↓ | | 0.5 |
|  | Creatinine | ≈ | | < 0.1 | ≈ | | < 0.1 | ↓ | | 0.2 |
| **Glycolysis and lactate production** | Glucose | ↓ | | 0.2 | **↑** | | 4.5 | **↑** | | 3.5 |
|  | Lactate | ↓ | | 3.2 | ↓ | | 0.2 | **↑** | | 1.9 |
|  | **Pyruvate** | ↓ | | 0.2 | **↑** | | 0.4 | **↑***** | | **1.4** |

Supplemental Table 3:

Metabolomics data changes overview in three mice groups checked against the control ND-WT mice. Non-significant changes (by t-test) not labeled. VIP (Variable Importance in Projection) score are depicted for each group.
Statistical significance: *p < 0.05, **p < 0.01, ***p < 0.001.

Supplemental Fig. 3

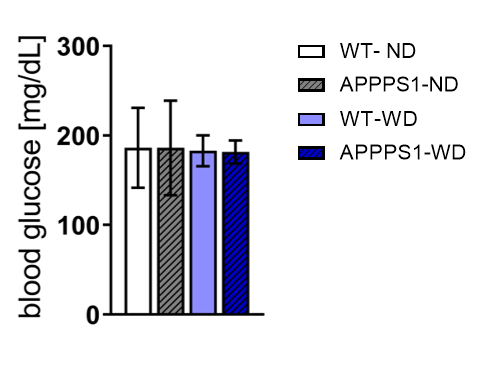

Supplemental Figure 3:

Mean blood glucose levels measured before [^18^F]FDG imaging for all 4 groups did not differ. WT-ND n = 11, APPPS1-ND n = 7, WT-WD n = 7, APPPS1-WD n= 8, post hoc Tukey corrected for multiple comparisons.

Supplemental Fig. 4

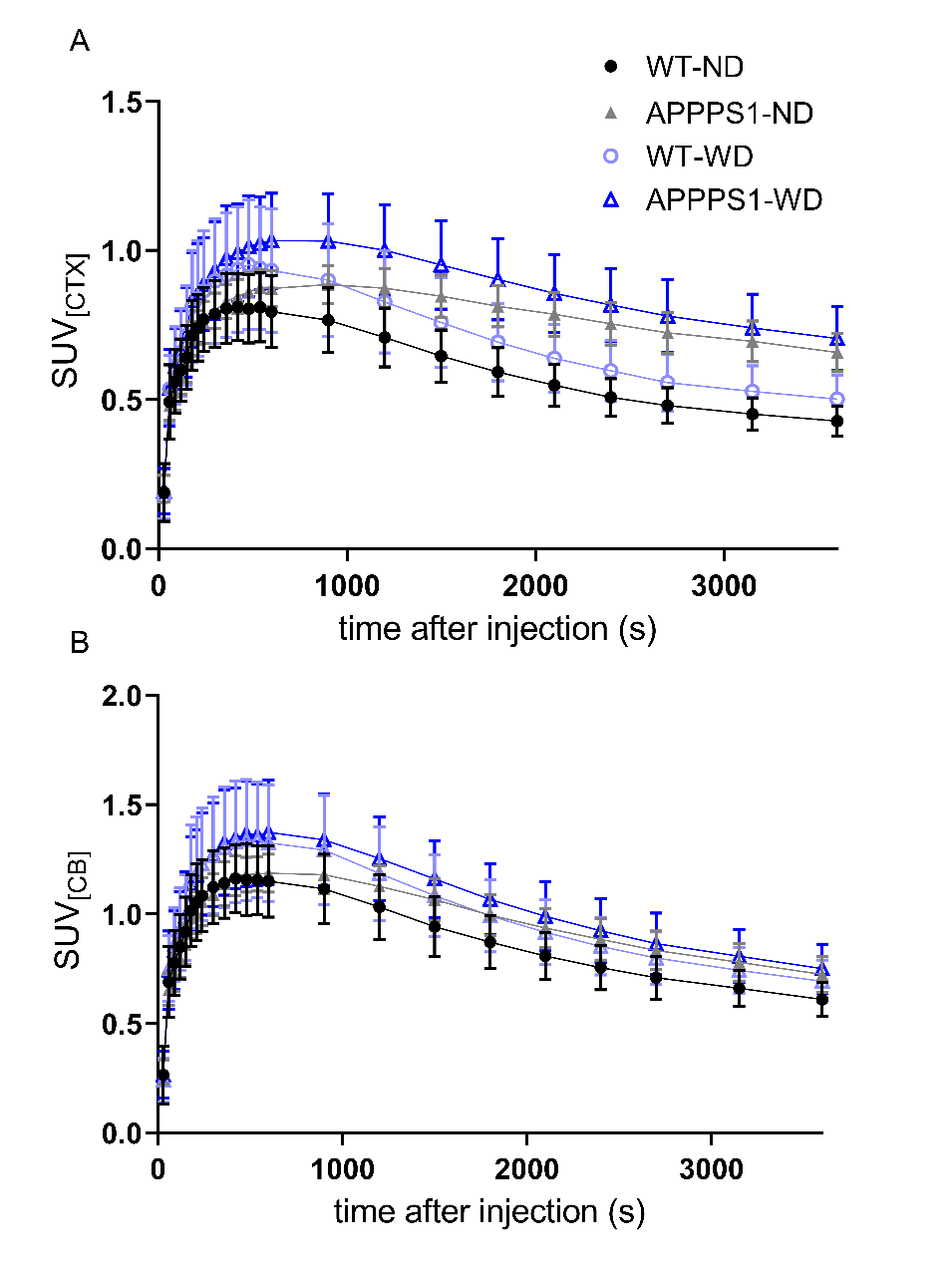

Supplemental Figure 4:

Time activity curves (TACs) showing mean ± SD [^18^F]GE-180 uptake in cortex (A) and cerebellum (B) over time for all 4 groups. WT animals depicted with circle, APPPS1 animals with triangles. ND-fed animals in black/grey, WD-fed animals in light/dark blue. WT-ND n = 8, APPPS1-ND n = 7, WT-WD n = 9, APPPS1-WD n= 10.

Supplemental Fig. 5

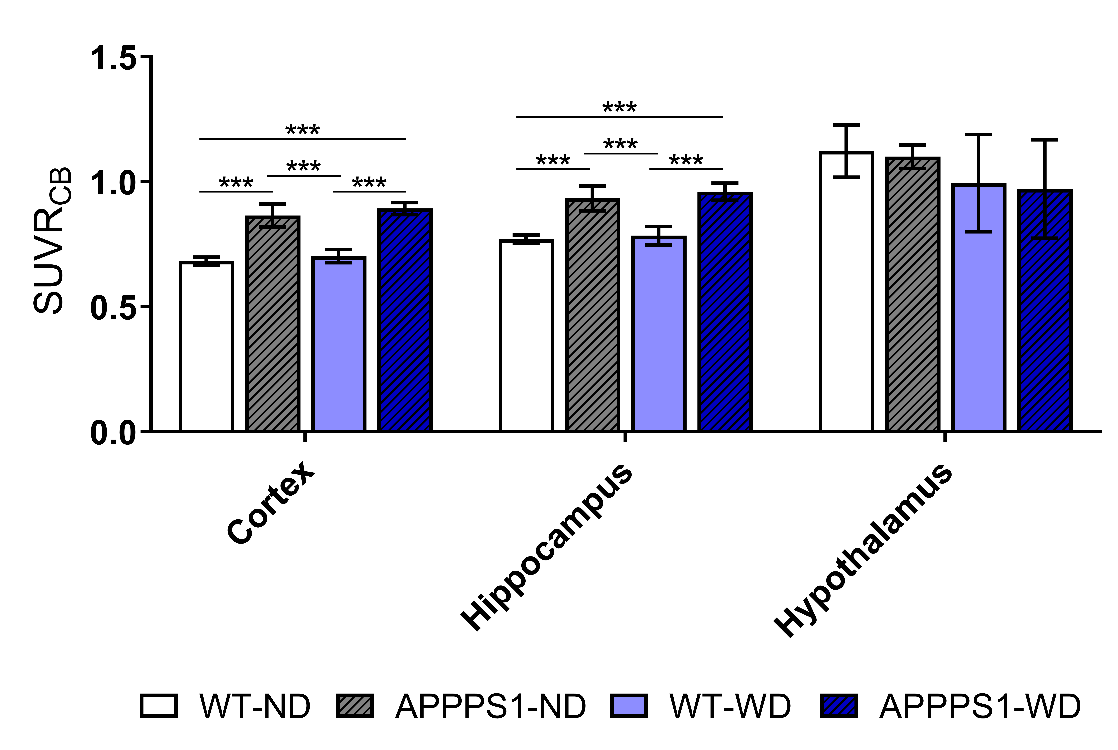

Supplemental Figure 5:

Mean ± SD SUV ratios (SUVR) for [^18^F]GE-180 in cortex, hippocampus and hypothalamus. The cerebellum was used as pseudo-reference region. Differences are observed between transgenic and wild-type animals in cortex and hippocampus independent of the diet. One-way ANOVA with multiple comparison analysis with post hoc Tukey correction. WT-ND n = 8, APPPS1-ND n = 7, WT-WD n = 9, APPPS1-WD n= 10. ***p<0.001.

Supplemental Fig. 6

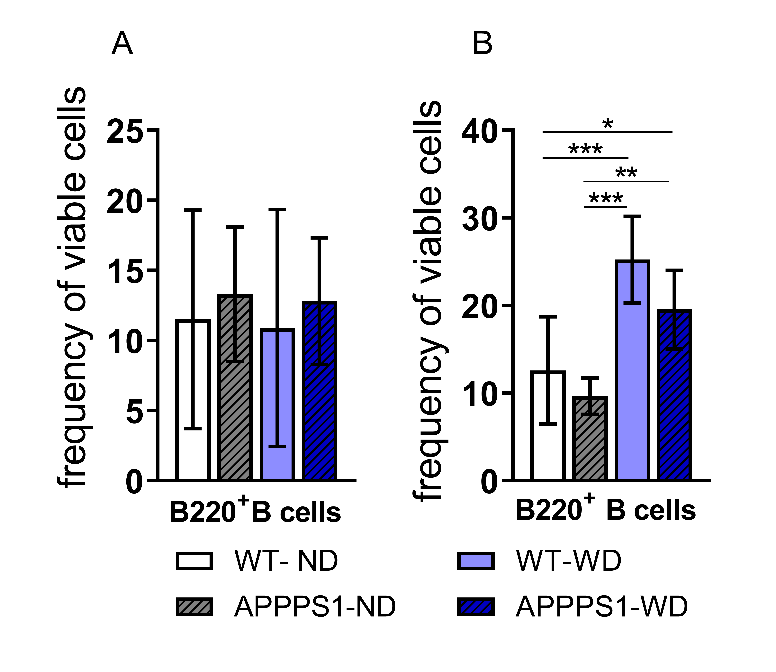

Supplemental Figure 6:

(A) Flow cytometry results showing no changed B cell populations in the brain, but (B) significantly elevated in WAT of WD-fed mice. Results in mean ± SD; *p<0.05, **p<0.01, ***p<0.001, post hoc Tukey corrected for multiple comparisons.

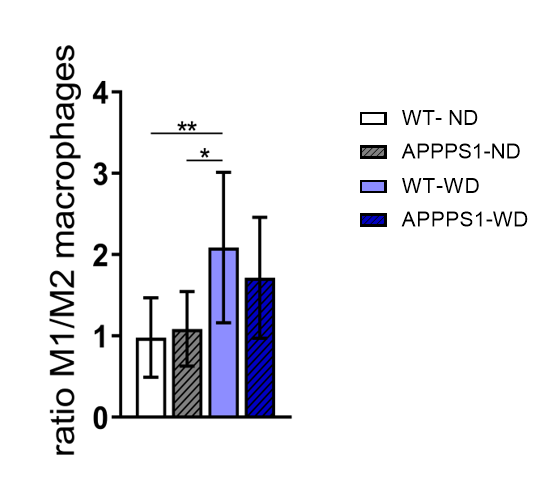
Supplemental Fig. 7

Supplemental Figure 7:

Flow cytometry results showing higher M1/M2 ratio in WD-WAT. *p<0.05, **p<0.01, post hoc Tukey corrected for multiple comparisons.

Supplemental Fig. 8

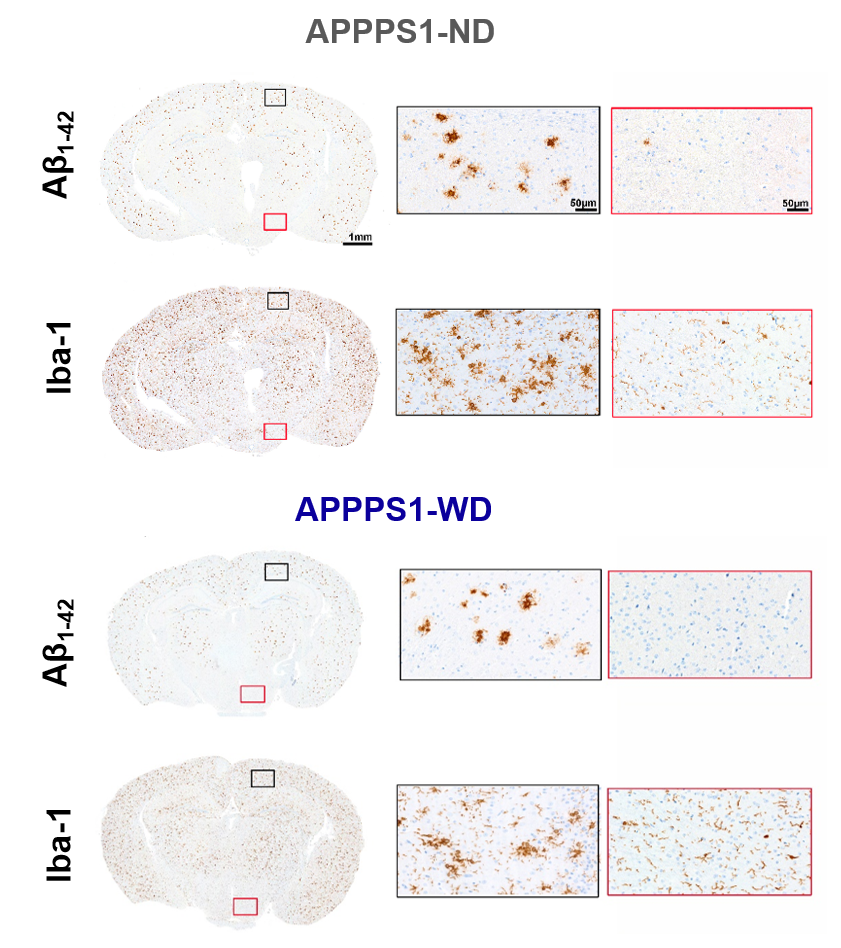

Supplemental Figure 8:

Comparison of Iba-1 reactivity in amyloid-rich (regions marked in black) and amyloid-poor (marked in red) regions of the APPPS1 brains showing that the microglia is highly activated in the regions of high plaque load. No differences between the diets are detected. Magnifications are depicted accordingly (Scale bar 50 µm). APPPS1-ND n = 3; APPPS1-WD n =3. Scale bar whole brain 1 mm.

Supplemental Fig. 9

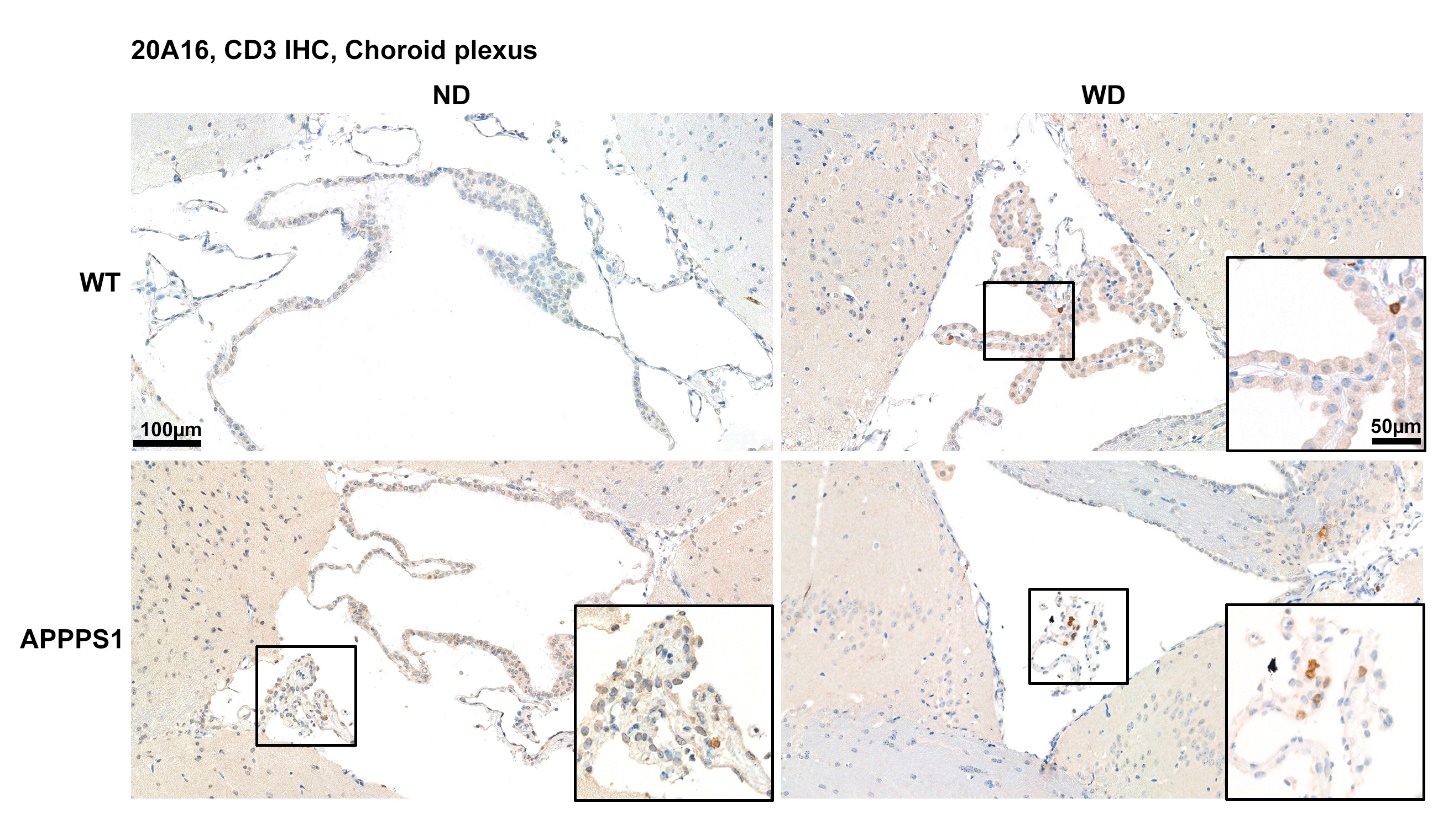

Supplemental Figure 9:

Exemplary images of CD3 + T cells infiltration in choroid plexus. Intragroup variability was high in APPPS1-WD animals, but showed tendency towards higher infiltrates than other groups. WT-ND n = 2; APPPS1-ND n = 3; WT-WD n = 3; APPPS1-WD n =3. Scale bar overview 100µm; magnification 50µm).
